# Cell-type-resolved *NRXN1* isoforms in human brain and hiPSC cortical organoids

**DOI:** 10.1101/2025.11.11.687875

**Authors:** Lei Cao, Yu Fan, Sadaf Ghorbani, Jessica Mariani, Yanchun Zhang, Michael B. Fernando, Jaroslav Bendl, John Fullard, Susana Isabel Ramos, Edward A. Mead, Nicola A.L. Hall, Gintaras Deikus, Kristin G Beaumont, Bohan Zhu, Sai Ma, Robert Sebra, Nadejda Tsankova, Panos Roussos, Kristen J. Brennand, Gang Fang

## Abstract

*NRXN1* undergoes extensive alternative splicing that generates a highly diverse repertoire of isoforms, diversifying protein–protein interactions, shaping synaptic specialization, and contributing to neuropsychiatric disease when disrupted. However, the splicing landscape of *NRXN1* across distinct human brain cell types remains poorly defined. Because *NRXN1* expression level is relatively low in adult brains and human induced pluripotent stem cell (hiPSC) derived neurons, single-cell long-read cDNA sequencing often provides insufficient coverage to capture its full isoform diversity, particularly across heterogeneous cell populations. To address this gap, we developed an integrative sequencing strategy that combines single-cell transcriptomics, targeted enrichment of *NRXN1* transcripts, and long-read sequencing. This approach reveals a comprehensive catalog of cell-type resolved *NRXN1* isoforms across adult and fetal human postmortem brains as well as hiPSC-derived cortical organoids. From the adult prefrontal cortex (PFC) region, our analyses reveal distinct splicing programs across interneuron subtypes, pyramidal neurons, and glial lineages. Comparisons between prenatal and adult brains indicate that *NRXN1* isoform profiles are established during early development and remain stable throughout neuronal maturation. In hiPSC organoid models, splicing profiles partially mirror both developmental and mature brain patterns, with a subset of splice sites exhibiting cell type – and stage-specific divergence. In the cerebellum of an autism case and in organoids derived from schizophrenia patients, carrying non-recurrent heterozygous *NRXN1* deletions, we identify disrupted isoform expression and mutant *NRXN1α* isoforms and characterize their distribution across diverse cell types. Using this integrated long-read sequencing framework in patient-derived organoids, we further show that antisense oligonucleotide (ASO) targeting of mutant splice junctions reduces gain-of-function *NRXN1* isoforms and reshapes splicing patterns in a cell type–specific manner, providing a platform for evaluating ASO efficiency. Together, these findings help understand the cell type–specific splicing landscape of *NRXN1* and establish a framework for decoding cell type-specific isoform diversity and assessing transcript-targeted therapies for genes with low expression levels.

## INTRODUCTION

Alternative splicing (AS) is a key post-transcriptional process in the human brain that generates diverse mRNA isoforms, contributing to the brain’s functional complexity^1–4^. Presynaptic cell-adhesion molecules known as neurexins (*NRXN1-3*) stand out as canonical examples of AS-driven synaptic diversity, playing central roles in synapse formation, specification, and plasticity^5–9^. The *NRXN1* gene generates an exceptional number of isoforms through six major alternative splice sites (SS1-6)^10–12^, which are essential for fine-tuning synaptic properties and diversification. For instance, alternative splicing at splice site 4 (SS4) is known to reshape ligand binding^13–16^, while differences at SS3 in hippocampal GABAergic interneurons, specifically parvalbumin-positive (*PV*+) and cholecystokinin-positive (*CCK*+) subtypes, have been linked to distinct presynaptic calcium channel usage and modulatory receptors^17,18^. Aberrant alternative splicing due to *NRXN1^+/−^* deletions disrupt the excitatory–inhibitory balance by weakening excitatory transmission in glutamatergic neurons and enhancing inhibitory transmission in GABAergic neurons^19^, underscoring how precise isoform regulation is critical for maintaining synaptic function.

A complete, cell-type-resolved isoform catalog of *NRXN1* remained elusive, particularly due to the technological limitations for detection of the large number of neurexin isoforms and their extensive cellular diversity in the human brain. Methods such as patch pipette-based qPCR^20^ and next generation sequencing^20^ can examine various insertions at alternatively spliced segments but do not assess the combinatorial nature of splicing events on a single mRNA molecule and lack the throughput to capture the *NRXN1* splicing patterns in large populations of diverse cells. Long-read platforms, coupled with single-cell RNA barcoding, represent a major development to study splicing of complex genes such as *NRXN1* in a cell type-specific manner. However, due to the low expression level of *NRXN1*, these transcriptomic approaches provide insufficient depth to comprehensively assess the large repertoire of *NRXN1* isoforms.

To overcome these technical barriers, we developed an integrative sequencing strategy that combines (1) probe-based long-read targeted-seq (long-read Capture-seq)^21,22^ with (2) single-cell/single-nucleus RNA isoform sequencing for in-depth *NRXN1* isoform profiling, complemented with (3) an optimized long-read RACE-seq ^23,24^ (Rapid Amplification of cDNA Ends) based enrichment module to focus on specific mutant isoforms of interest. In the core idea of this integrative strategy, single-cell transcriptomics is used to provide cell-type deconvolution, while targeted capture is used to enrich *NRXN1* transcripts for in-depth analysis; long read sequencing is then employed to achieve the crucial full-length isoform resolution. The use of both probe-based targeting and RACE-seq (PCR-based) enrichment complement each other in recovering the diversity of *NRXN1* isoforms (the strength of probes) and detecting rare or low-abundance mutant isoforms (the strength of RACE-seq). Using this integrative strategy, we mapped a comprehensive catalog of cell-type-specific *NRXN1* isoforms across diverse neuronal and glial populations. From the PFC region of the human adult brain, we revealed distinct splicing profiles across GABAergic interneuron, pyramidal neurons and glial lineages. Interneurons derived from the same developmental origin exhibited similar AS patterns, whereas those derived from different origins showed divergent profiles. Analyses of the AS pattern in fetal brain suggest that *NRXN1* splicing profiles are defined in early development and remain largely consistent during neural maturation. Using hiPSC cortical organoids derived from patients with *NRXN1^+/−^* deletions and controls, we observed that glial progenitor cells and upper-layer excitatory neurons (Ex-L2/3) are enriched for the mutant isoforms, and we found that the organoids partially recapitulate the *NRXN1* splicing patterns of both developing and adult brains. In the postmortem brain tissue from an autistic patient case carrying a heterozygous *NRXN1* deletion, we found that the mutant isoforms transcribed from deletion allele were enriched in molecular layer interneurons (MLI1/2) and astroglia cells. These findings provide insights into the complex alternative splicing of *NRXN1* across brain cell types, along with a generally applicable framework for decoding cell type-specific isoform diversity for important genes with low gene expression levels.

## RESULTS

### An integrated strategy reveals a cell-type resolved catalog of *NRXN1* isoforms across human brain and hiPSC cortical organoids

Single-cell long-read transcriptomics provides a unique ability to resolve full-length *NRXN1* isoforms across cell types, but whole transcriptomic profiling does not provide sufficient depth of *NRXN1* due to its very low expression level. From the published human postmortem brain single-nuclei Iso-seq data^25^, *NRXN1* reads constituted only 0.09-0.2% of the total reads across different cell types, underscoring the need for a targeted enrichment strategy (**Fig. 1a, upper panel**). To maximize isoform recovery while preserving cell-type resolution, we used a targeted sequencing strategy with two complementary components (**Fig. 1b**). To generate a comprehensive isoform catalog, we designed capture probes tiled across all *NRXN1* exons at 1× density, which achieved a ∼36-96-fold enrichment of *NRXN1* reads over the non-targeted sequencing method (**Fig. 1a, lower panel**). We built on Tequila-seq, which uses isothermal amplification of capture probes from low-cost oligonucleotide templates^21^ to reduce per-reaction costs by two to three orders of magnitude while maintaining accurate transcript quantification across gene panels of varying sizes. To recover cell-type specificity for isoforms of interest, particularly the mutant isoforms transcribed from the allele carrying exonic deletions, we adapted the RACE-seq (Rapid Amplification of cDNA Ends) method from short-read and optimized for long-read sequencing to achieve a more sensitive detection of low-abundance isoforms. To link isoforms to specific cell types, we matched the UMIs and cell barcodes on the *NRXN1* reads (from *NRXN1*-targeted long-read sequencing data) with the UMIs from the cell cluster information by short-read single-cell RNA-seq data (**Methods**). This design leveraged the sequencing depth and accuracy of short reads together with full-length isoform resolution by long reads, enabling precise assignment of complex *NRXN1* isoforms to distinct cell types.

**Figure 1.**
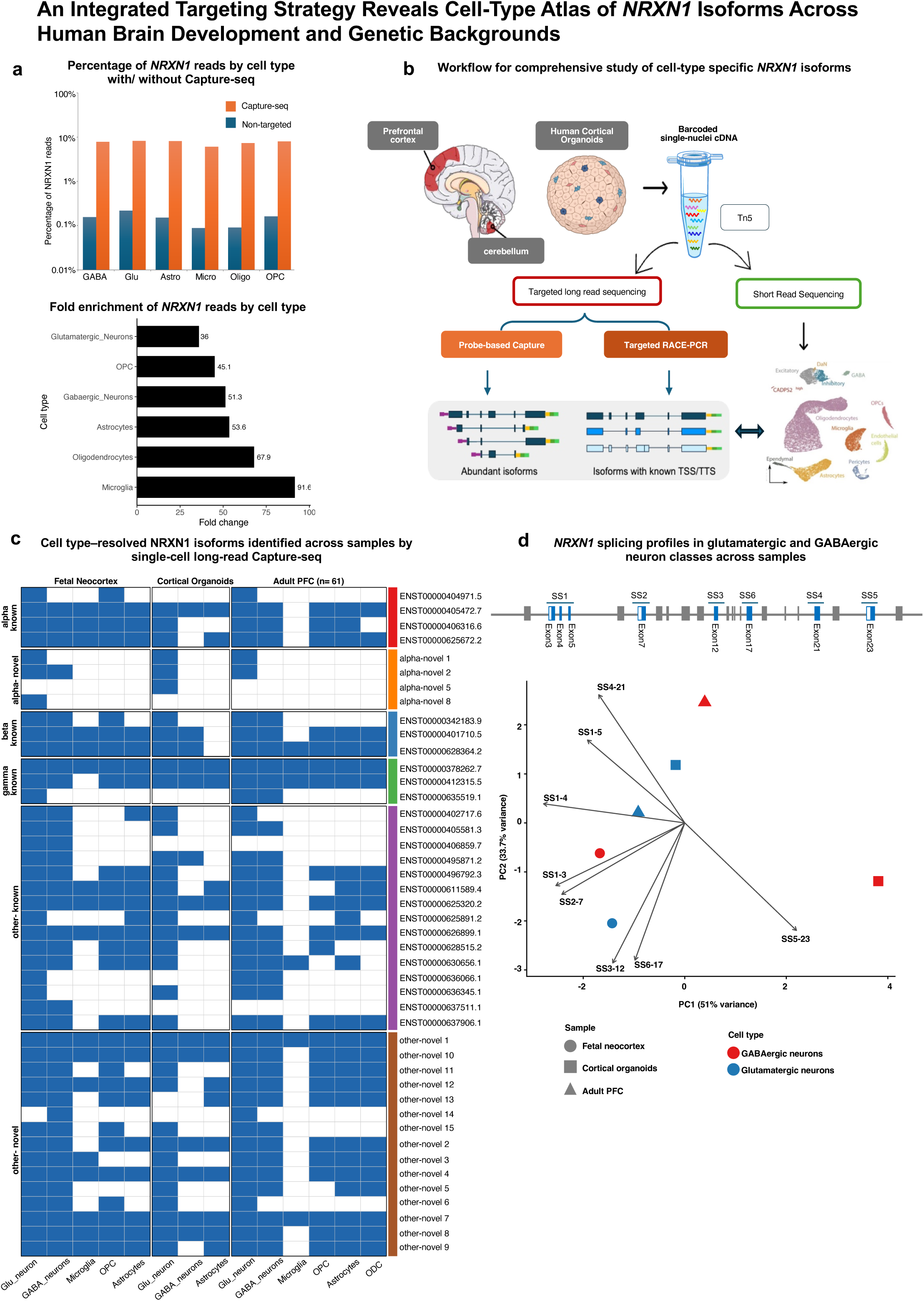
A targeting strategy reveals cell-type catalog of *NRXN1* isoforms. (a) *Upper panel* Bar graph showing the percentage of *NRXN1* reads detected by long-read Capture-seq (orange) and non-targeted iso-seq (blue) across major brain cell classes. GABA, GABAergic neurons; Glu, glutamatergic neurons; Astro, astrocytes; Micro, microglia; Oligo, oligodendrocytes; OPC, oligodendrocyte progenitor cells. *Lower panel:* Fold change of *NRXN1*-mapped reads obtained by long-read Capture-seq relative to the non-targeted single-cell Iso-seq. (b) Schematic illustrating the comprehensive characterization of cell-type-specific *NRXN1* isoforms using the targeting strategy. Full-length single-cell/nuclei cDNA is divided into two parts: one processed for a short-read library using Tn5 transposase and sequenced to identify cell types; the other subjected to probe-based long-read Capture-seq and RACE-seq to detect *NRXN1* isoforms with matched barcodes and UMIs (see Methods). (c) *NRXN1* isoforms identified by single-cell long-read Capture-seq across samples. Each row represents a unique isoform, categorized as α, α-mutant, β, γ, other known, or novel. Ensembl-annotated isoforms are labeled with ‘ENST’ IDs. The heatmap shows presence (blue) or absence (white) across adult prefrontal cortex, fetal neocortex, and human cortical organoids. (d) PCA biplot of NRXN1 splicing profiles in GABAergic and glutamatergic neurons across samples. Principal component analysis of mean inclusion fractions for exons at 6 splicing sites (SS1-6). Arrows indicate SS event loadings with arrow direction showing which sample groups have higher inclusion of corresponding events. The arrow length reflects the magnitude of contribution to variance. A schematic of *NRXN1* gene structure is shown, with splice sites (SS1–SS6) and alternatively spliced exons at each site highlighted in blue.

We applied this workflow to single-cell or single-nucleus cDNA from multiple sources, including the PFC of adult brains, gestational cortical samples and cerebellar tissue from an *NRXN1*^+/–^ autism patient. In addition, cortical organoids derived from schizophrenia patients harboring unique *NRXN1*^+/–^ deletions were profiled to assess the impact of these variants on splicing regulation. After isoform identification and quality filtering, using long-read Capture-seq, we identified 50 distinct *NRXN1* isoforms across all datasets, including 27 previously annotated (five α, five β, five γ, and twelve known others) and 23 novel isoforms not represented in Ensembl^26^ (**Extended Fig. 1a**). Cell barcode matching revealed 25 annotated and 19 novel isoforms expressed in glutamatergic and GABAergic neurons across samples (**Fig. 1c**). Expression of these novel isoforms was validated across our in-house datasets and confirmed using independent long-read transcriptomes from postmortem human brains reported by others^25,27^ using miniQuant-based quantitative analysis^28^ (**Extended Fig. 1b**). Among all the *NRXN1* isoforms, **alpha** isoforms constitute the largest and most structurally complex class. We applied long-read RACE-seq, a more sensitive method to single-cell or single-nucleus cDNA of all samples and identified 103 **alpha** isoforms including 57 that have been reported previously (**Extended Fig. 1c**). We compared exon inclusion across *NRXN1* splice sites (SS1–SS6) in GABAergic and excitatory neurons across all samples and performed PCA. Exons 3 and 4 at SS1 and exon 7 at SS2 show consistently higher inclusion in glutamatergic neurons, whereas exon 21 at SS4 is enriched in GABAergic neurons in adult PFC. Notably, GABAergic neurons in cortical organoids display a distinct splicing profile driven by exon 23 at SS5 (**Fig. 1d)**.

### *NRXN1* exhibits distinct splicing patterns across cell types in adult human brains

To understand *NRXN1* alternative splicing patterns across mature neurons in the human brain, we applied the probe-based long-read Capture-seq on the single-nucleus cDNA from the PFC region of 61 adult donors^27^ (**Fig. 2a, b; Extended Fig. 2**). We generated 2.43M *NRXN1* reads, among which 1.13M had 10x single cell barcodes, and 1.03M reads sharing the barcodes with short-read data. By integrating the barcode information from short– and long-read data, a total of 125,199 cells were identified across all donors. From the long-read Capture-seq of the single nuclei cDNA pools, we identified 46 isoforms with confident cell barcodes and UMIs matched to short-read data, including 6 alpha isoforms not annotated but reported previously, and 14 isoforms not annotated in Ensembl^26^ but later validated by our in-house brain dataset (**Fig. 1c, Extended Fig. 1**).

**Figure 2.**
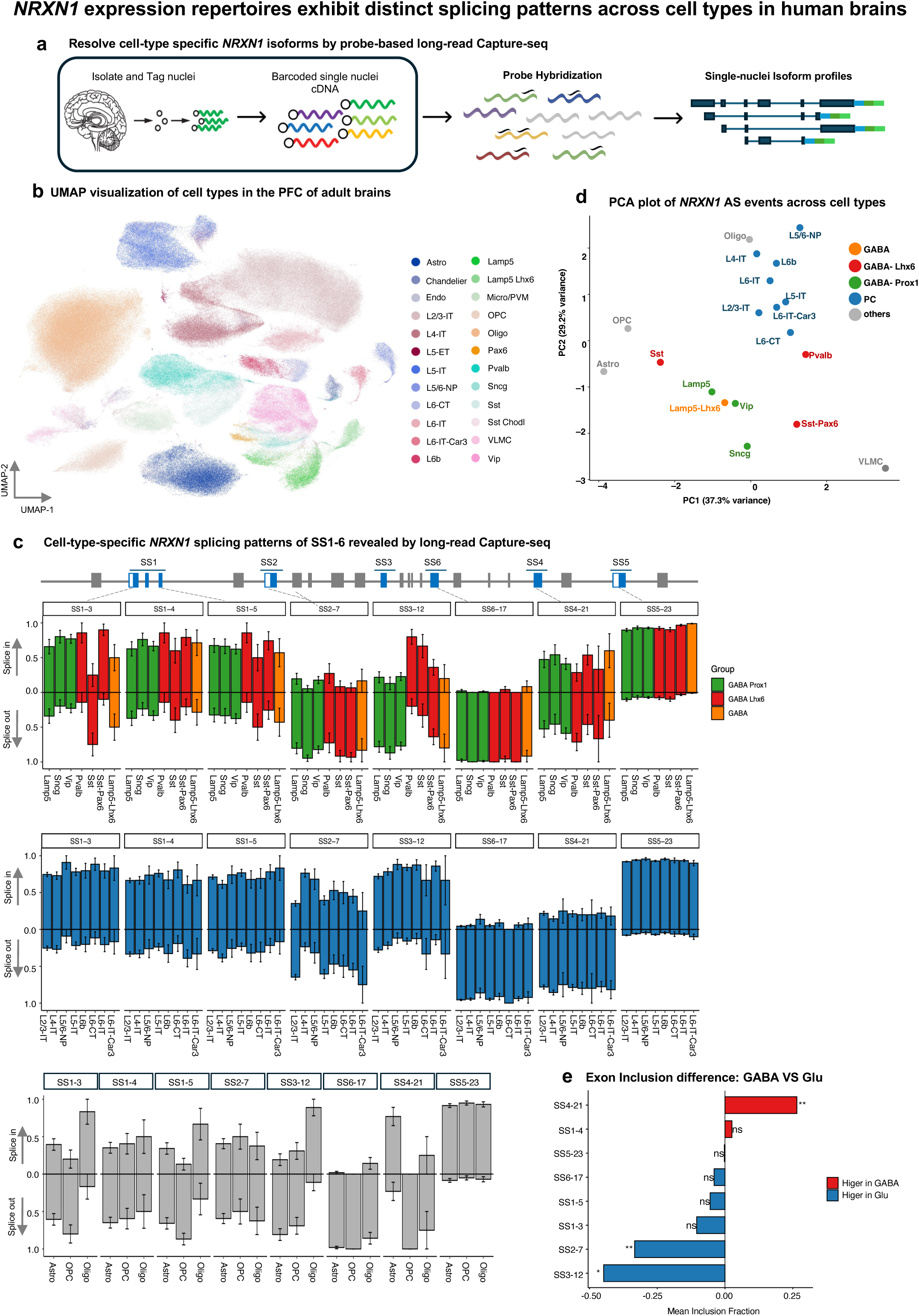
*NRXN1* expression repertoires exhibit distinct splicing patterns across cell types in human brains. (a) Workflow overview of long-read Capture-seq applied to single-nucleus cDNA from adult PFC. (b) UMAP visualization of putative cell types in the adult PFC. Colored clusters represent neuronal and glial subtypes. (c) Normalized *NRXN1* splicing events at six splicing sites (SS1–SS6) in GABAergic interneurons (top), glutamatergic excitatory neurons (middle), and glial lineages (bottom). SS1–3 represents exon 3 inclusion/ exclusion at splicing site, similar for SS1-4, SS2-7, *etc.* Each bar represents a cell class. Within GABAergic interneurons, *PROX1^+^-*INs are colored in green, *LHX6^+^-*INs in red, and *LAMP5⁺ LHX6^+-^*INs in orange. Exon inclusion (“splice-in”) is shown upward and exclusion (“splice-out”) downward. Data represent mean ± SEM. A schematic of *NRXN1* gene structure is shown, with splice sites (SS1–SS6) and alternatively spliced exons at each site highlighted in blue. (d) PCA plot based on *NRXN1* splicing patterns across splice sites SS1–SS6, showing that *PROX1⁺* –INs, *LHX6⁺* –INs, and PCs form distinct, separately clustered classes. Colors correspond to cell types as in (c): *PROX1⁺-* INs (green), *LHX6⁺*-INs (red), *LAMP5⁺ LHX6^+^*-GABAergic neurons (orange), glutamatergic neurons (blue), and glial lineages (gray). (e) The difference in mean exon inclusion fraction between GABA and Glu neuronal cell types (Δ = mean − mean) at 6 splicing sites. Positive values indicate higher inclusion in GABA neurons; negative values indicate higher inclusion in Glu neurons. Significance labels denote BH-adjusted *p*-values from Welch’s t-tests (** *p* < 0.01; * *p* < 0.05; ns, not significant).

The brain comprises a great diversity of cell types, and studying neurons with shared developmental markers offers a means to uncover how *NRXN1* alternative splicing contributes to this cellular and functional diversity. Based on developmental markers^27^, neurons were categorized as Neurod2-expressing pyramidal cells^28^ (PC), *LHX6*-expressing interneurons^29^ (*LHX6*^+^-INs originated from medial ganglionic eminence, MGE), *PROX1*-expressing interneurons^30^ (*PROX1*^+^-INs originated from caudal ganglionic eminence, CGE), and *LAMP5*^+^ *LHX6^+^* GABAergic neurons (*PROX1^+^* labeled as *LAMP5*). *LHX6*^+^-INs can be further subclassified into *PVALB^+^*and *SST^+^* INs, while *PROX1*^+^-INs are divided into *LAMP5^+^*, *SNCG*^+^, and *VIP*^+^ INs. GABAergic INs are a highly heterogeneous collection of cell types, characterized by their unique spatial and temporal capabilities to influence neuronal circuits^31–33^. We observed that GABAergic interneurons showed the greatest *NRXN1* splicing diversity (**Fig. 2c, top**). Splicing patterns were more variable at SS1, SS3, and SS4, whereas SS2 and SS6 were invariably excluded and SS5 was consistently included. Within the *PROX1^+^*-INs, *the LAMP5^+^, SNCG*^+^ and *VIP^+^* subtypes share similar patterns across all 6 alternative splicing sites, while *PVALB^+^* and *SST^+^*, subclasses of *LHX6^+^*-INs, differed at SS1. We also identified a *LAMP5^+^ LHX6^+^* subclass that shared splicing patterns with the *PROX1^+^*-INs and *SST^+^* of *LHX6^+^*– INs (**Fig. 2c**). In PC neurons (**Fig. 2c, middle**), we found: (1) across cortical layers, exons at SS1, SS3 and SS5 were dominantly spliced in (“in”; upward bars), while exons at ASS6 and SS4 are dominantly spliced out; (2) layer-specific differences at SS2, where exon 7 was often spliced out in PCs of layers L2/3-IT, L5-IT, and L6-IT-Car3, but spliced-in into most PCs of layers L4-IT, L5/6-NP, L6B, L6-CT, and L6-IT. Among glial cells (astrocytes, oligodendrocyte progenitor cells (OPCs), oligodendrocytes), exon 17 at SS6 was consistently spliced out, whereas exon 23 at SS5 was spliced in. Oligodendrocytes showed spliced-in for exon1 and exon5 at SS1, which were dominantly spliced-out in astrocytes and OPCs. At ASS4, exon 21 was included in astrocytes but not in OPCs or oligodendrocytes, consistent with prior studies showing that astrocytic neurexins splice SS4 at high ratios compared to excitatory neuronal neurexins^34,35^.

Building on the striking patterns of *NRXN1* alternative splicing observed in the human brain, we conducted principal component analysis (PCA) of splicing levels across cell types, which separately clustered *PROX1^+^*-INs, *LHX6^+^*-INs and PCs classes (**Fig. 2d**). To identify *NRXN1* splicing events distinguishing glutamatergic (Glu; n = 9) and GABAergic (GABA; n = 4) neurons, we compared mean exon inclusion across 6 splice-site events using Welch’s t-tests with Benjamini–Hochberg correction. SS2-7 and SS3-12 exhibited higher inclusion in Glu neurons, whereas SS4-21 was preferentially included in GABA neurons (**Fig. 2e**). Overall, these findings suggest that *NRXN1* splicing profiles exhibit substantial differences across various neuronal populations and glial lineages. Within GABAergic neurons, interneurons derived from the CGE (*PROX1^+^*-INs subtypes) are closely clustered by the splicing patterns, yet different from INs originating from the MGE (*LHX6^+^*-INs subtypes).

### *NRXN1* splicing profiles are conserved between fetal and adult neurons within defined neurogenic lineages

To determine whether the *NRXN1* splicing patterns of the adult brain are defined as early as in the human developing neocortex, we generated a snRNA-seq dataset from two unfixed snap-frozen postmortem neocortex samples obtained from the second and third trimesters of gestation (24 and 33 gestational weeks, gw). After quality control, we obtained the transcriptomic profile of 52,426 nuclei, with a median of 2,910 genes, 6,156 UMIs from each nucleus. We identified 10 primary cell types from the single cell transcriptome: five neuronal clusters, including excitatory neuronal subpopulations expressing layer-specific cortical marker genes (EX-L2/3, EX-L4/5/6); inhibitory neuronal subpopulations (IN-CGE, IN-MGE and IN-Reelin); and glial cell populations, radial glia and astrocyte (RG/AC) clusters, astrocytes, OPCs, transient glial intermediate progenitor cell (gIPC) population, microglia and *FLT1*+ blood vessel cells (BVCs)^36^ (**Fig. 3a**). We further validated our cell annotations by comparing top differentially expressed genes (DEGs) per cell cluster to published canonical lineage and proliferation markers^36,37^ (**Fig. 3c, Supplementary Table 3**). Next, we analyzed how cell type proportion varied between prenatal neocortex and adult PFC. The 33 gw neocortex sample had more glial, INs-MGE and INs-CGE, whereas the 23 gw sample had more INs-Reelin and excitatory neurons EX-L4/5/6 (**Fig. 3b**). Neuronal populations constituted most cells in the fetal neocortex, whereas glial transient intermediate progenitor cells were exclusively present in fetal neocortex and subsequently disappeared or transformed along with the cortical development in adult PFC, which is consistent with the stages of neurogenesis in human cerebral cortex^38,39^ (**Extended Fig. 3**).

**Figure 3.**
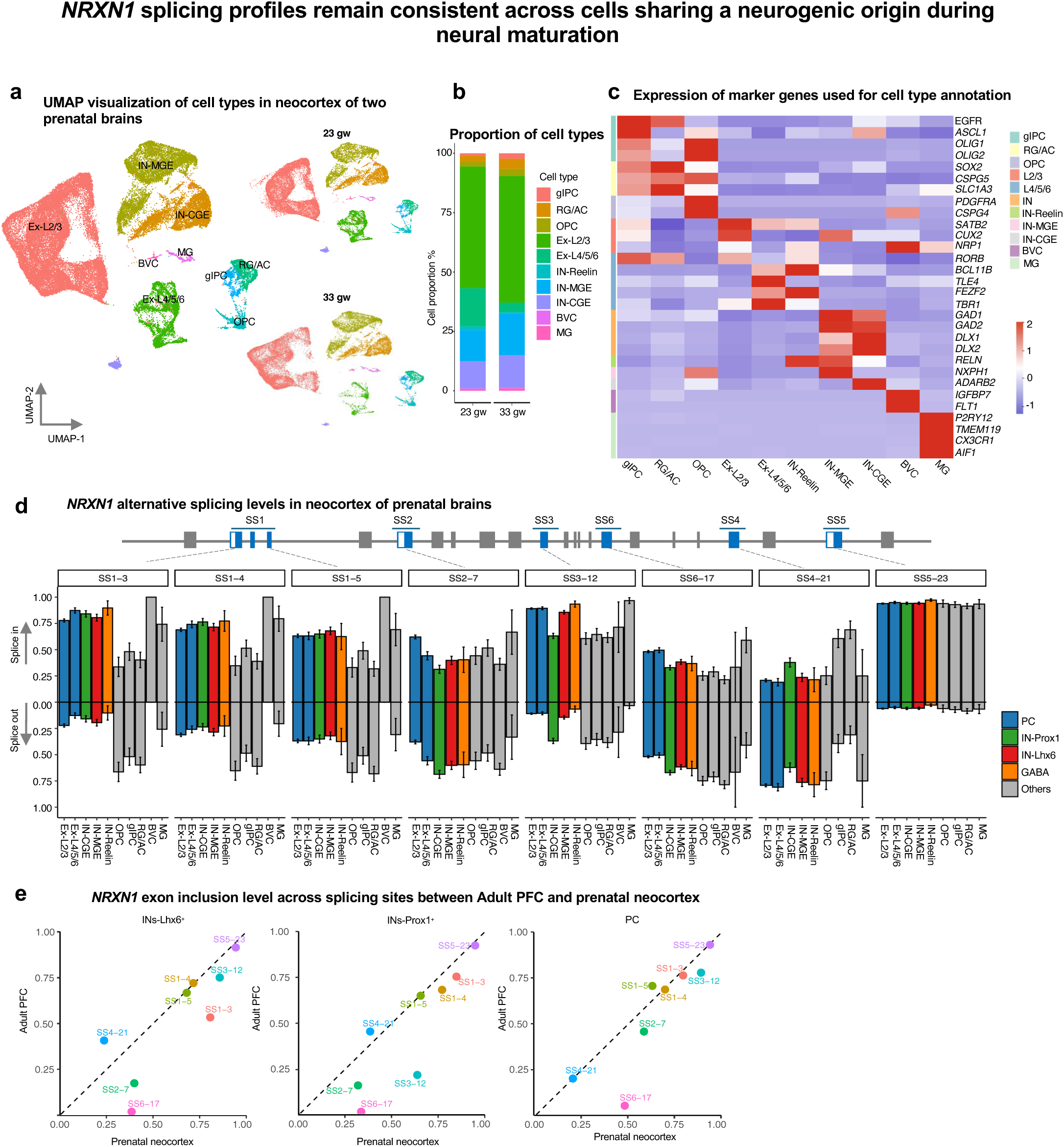
*NRXN1* splicing profiles remain consistent across cells sharing a neurogenic origin during neural maturation. (a) UMAP visualization of putative cell types in the neocortex from two fetal brains. *Left:* merged UMAP of combined datasets; *Right:* separate UMAPs showing neocortex samples from 23 gw and 33 gw. Colored clusters represent neuronal and glial subtypes in the fetal neocortex. Abbreviations: gIPC, glutamatergic intermediate progenitor cells; RG/AC, radial glia/astrocyte lineage; Ex, excitatory neurons; IN, inhibitory neurons; BVC, blood vessel cells; MG, microglia. (b) The proportions of each cell class in 23 gw and 33 gw neocortex samples. Colored blocks indicate neuronal and glial subtypes. (c) Heatmap of marker genes expression used for cell-type annotation. Each row represents a marker gene and each column a cell class; darker red indicates higher expression. Colored blocks indicate neuronal and glial subtypes in fetal neocortex. (d) Normalized *NRXN1* splicing events at SS1–SS6 in *PROX1*⁺-INs (green), *LHX6*⁺-INs (red), glutamatergic excitatory neurons (blue), and glial lineages (gray). A schematic of *NRXN1* gene structure is shown, with splice sites (SS1–SS6) and alternatively spliced exons at each site highlighted in blue. Exon inclusion (“splice-in”) is shown upward and exclusion (“splice-out”) downward. Data represent mean ± SEM. (e) Pearson’s correlation of *NRXN1* exon inclusion levels across splicing sites (SS1–SS6) between adult PFC and prenatal neocortex in *LHX6*⁺-INs (left), *PROX1*⁺-INs(middle), and PC neurons (right).

To determine *NRXN1* splicing patterns by cell type in the fetal neocortex, we applied the probe-based long-read Capture-seq on the single nuclei cDNA. Based on regional marker genes, exon 23 at SS5 was predominantly spliced-in across all cell types, whereas exons at other splice sites showed variable inclusion. In most (>60%) PCs across all layers, exons at SS1 and SS3 were consistently spliced in, exon 21 at SS4 was spliced out, and exon 17 at SS6 showed variable usage. In contrast, exon 7 at SS2 was differentially spliced between upper-layer (L2/3) and deeper-layer (L4/5/6) PCs. Similarly, in GABAergic interneurons (INs), exons at SS1 and SS3 were largely spliced in, while exons at SS2, SS4, and SS6 were spliced out. Notably, the proportion of IN-*LHX6⁺* cells was higher than that of IN-*PROX1*⁺ cells at SS3 (spliced in) and SS4 (spliced out). The glial cells were variably spliced at SS1, SS2, SS6 and SS4, but consistently spliced-in at SS3 and SS5 (**Fig. 3d**). We next compared *NRXN1* splicing in matched neuronal populations between prenatal neocortex and adult PFC. Exon inclusion level across 6 splicing sites were highly correlated between prenatal neocortex and adult PFC for *LHX6*^+^-INs (*Pearson’s r* = 0.82, *p* = 0.01), *PROX1*^+^-INs (*Pearson’s r* = 0.87, *p* = 0.005), PCs (*Pearson’s r* = 0.87, *p* = 0.005) and glial populations (OPC, *Pearson’s r* = 0.9, *p* = 0.003; RG/AC, *Pearson’s r* = 0.82, *p* = 0.01) (**Fig. 3e, Extended Fig. 3**). These findings suggest that *NRXN1* splicing profiles are defined in early development and remain largely consistent during neural maturation.

### *NRXN1⁺/⁻* splicing profiles in human cortical organoids recapitulate developmental and mature features and identify cell types expressing the mutant isoforms

From hiPSC-derived two-dimensional (2D) neurons with a 3′-*NRXN1* deletion, we previously uncovered dozens of mutant isoforms exclusively from the deletion-carrying allele (mainly affecting alpha and beta isoforms)^12^. Later, we demonstrated that this exonic deletions at the 3’ site disrupted normal *NRXN1* splicing and isoform expression, leading to distinct phenotypes in induced glutamatergic (iGLUTs) and GABAergic (iGABAs) neurons, primarily reflected in altered synaptic frequency^19,40^. Building on these observations, we sought to explore how the deletions affect the splicing events across cell types in a more complex neurodevelopmental system. We generated human cortical organoids (hCOs) from a control line and schizophrenia probands with the 3’deletion (3’-del) in parallel. hCOs were guided toward the forebrain^41^ (**Methods, Extended Fig. 4**). To profile cell types in the hCOs, we performed single cell RNA-seq on hCOs from two 3’-del cases (line 581 and line 641) and one control line (line 690) at day 150 with the 10x NextGEM 5pv2 kit, when neural activity is well characterized^42–44^. After quality control data processing steps, we merged high-quality scRNA-seq data from 3 cortical organoid samples into a core dataset of 27,325 cells. Following unsupervised clustering, we identified and annotated major cell types using an extensive curated list of known marker genes^19,41^, which consisted of transit-amplifying cell (TAC), neuronal intermediate progenitor cells (nIPCs) giving rise to distinct subpopulations of glutamatergic excitatory neurons (Ex-L2/3, Ex-L4/5/6) and GABAergic inhibitory neurons (INs, IN-*SST*,IN-Reelin); gIPCs leading to the subpopulations of radial glia and astrocyte clusters (RG/AC); as well as BVCs (**Fig. 4a**). We further validated our cell annotations by comparing gene expression signatures of our scRNAseq cell clusters to published brain organoid datasets as reference (**Fig. 4b, Supplementary Table 4**). We then compared cortical organoids derived from the control hiPSC line with the prenatal neocortex in terms of cell-type composition. We found gIPCs (OR = 2.06, *Fisher’s test*) and Ex-L2/3 (OR = 1.67, *Fisher’s test*) were significantly enriched in hCOs from the 3’-del lines than in control lines, while cell number of IN-Reelin were significantly low in 3’-del lines (OR = 0.03, *Fisher’s test*) (**Fig. 4c**).

**Figure 4.**
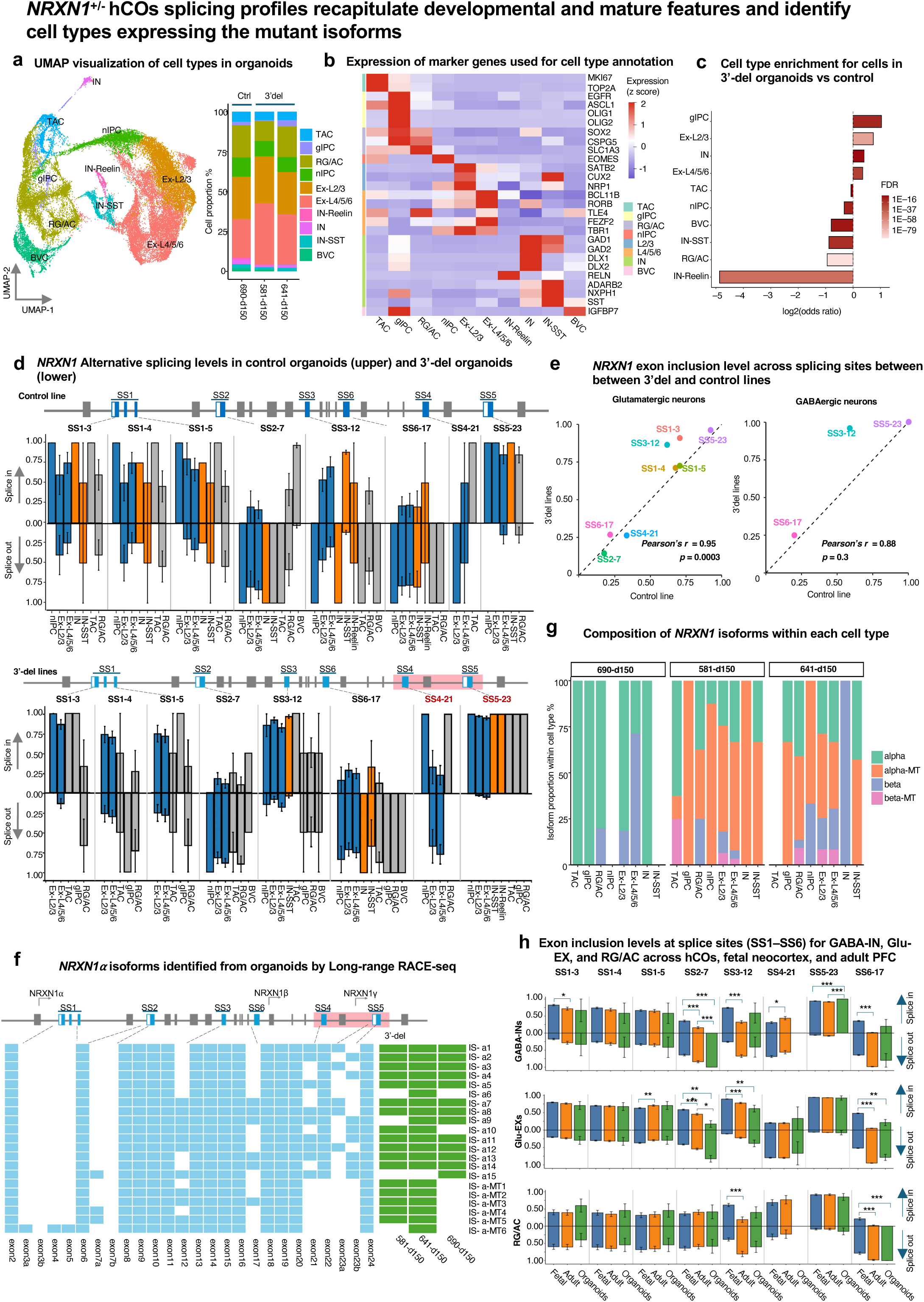
*NRXN1⁺/⁻* human cortical organoids (hCOs) splicing profiles recapitulate developmental and mature features and identify cell types expressing the mutant isoforms. (a) *Left:* UMAP of putative cell types in hCOs, with colored clusters representing neuronal and glial subtypes. *Right:* Proportions of each cell class in organoids derived from control line 690 or 3′-del lines (581 and 641). Abbreviations: gIPC, glutamatergic intermediate progenitor cells; nIPC, neuronal intermediate progenitor cells; TAC, transient amplifying cells; RG/AC, radial glia/astrocyte lineage; Ex, excitatory neurons; IN, inhibitory neurons; BVC, blood vessel cells. (b) Heatmap of marker gene expression used for cell-type annotation. Each row represents a marker gene and each column a cell type; darker red indicates higher expression. (c) Bar graph showing enrichment of specific cell types in 3′-del organoids compared with control lines. (d) Normalized *NRXN1* splicing events at SS1–SS6 in all cell classes from control hCOs (upper) and two 3′-del hCOs (lower). Each splicing graph is accompanied by a schematic of *NRXN1* gene structure with red shading indicating 3′-del–associated regions; alternatively spliced exons at each splice site are highlighted in blue. Exon inclusion is shown upward (“splice-in”), exclusion downward (“splice-out”). Data represent mean ± SEM. (e) Pearson’s correlation of *NRXN1* exon inclusion levels across splicing sites (SS1–SS6) between 3′-del and control hCOs. (f) *Left:* Schematic of *NRXN1α* isoform structures identified by long-read RACE-seq, along with a schematic of *NRXN1* gene structure. Red shading indicates regions associated with the 3′ deletion, and alternatively spliced exons at each splice site are highlighted in blue. Each row represents a unique *NRXN1α* isoform, with blue exons indicating inclusion and blanks indicating exclusion. *Right:* Presence or absence of each *NRXN1α* isoform across 3′-del and control hCOs (green = present; blank = absent). (g) The composition of *NRXN1* isoforms identified by long-read RACE-seq within each cell type across 3′-del and the control hCOs. (h) Exon inclusion levels at SS1–SS6 in GABAergic INs (top), glutamatergic excitatory neurons (middle), and RG/AC (bottom) across hCOs (blue), fetal neocortex (orange), and adult PFC (green). Statistical comparisons were performed using one-way ANOVA (*, *P* < 0.05; **, *P* < 0.01; ***, *P* < 0.001).Data represent mean ± SEM.

To better understand the greater splicing patterns of *NRXN1* across cortical organoid cell types, we applied the long-read Capture-seq on the full-length single-cell cDNA. From the control lines and those with the 3’-deletion, we generated 317,521 *NRXN1* reads, of which 144,450 with 10x single cell barcodes and 120,873 reads sharing the barcodes with short-read data. Integrating the barcode information across short– and long-read data, we identified 23,187 cells from all cortical organoid samples (**Extended Fig. 4**). Using these targeted single cell data, we first analyzed the *NRXN1* AS patterns in excitatory neurons, inhibitory neurons and non-neuronal cells for both 3’-del lines and control lines at the advanced maturation stage. Within the glutamatergic excitatory neurons (the major population in hCOs), *NRXN1* splicing patterns were highly similar between the control line and 3’del lines, except for differences in exon inclusion level at SS1 (exon 3) and SS3 (exon 12) (**Fig. 4d,e**). The ∼136-kb deletion impacting SS4 and SS5 (exons 21-23)^12,19^ slightly reduced exon inclusion at SS4 in the 3’-del lines (**Fig. 4e**).

As the deletion in the 3’-region of *NRXN1* generates mutant alpha and mutant beta isoforms, we sought to dissect which cell types are enriched for these mutant isoforms. As some of the mutant isoforms are low in abundance and the long-read Capture-seq works better with abundant isoforms, we employed long-read 5’RACE-seq to resolve alpha isoforms with single cell barcode information from the 5’GEX single cell cDNA (Methods). We identified 21 α-isoforms across all cortical organoids, of which 18 were not detected by long-read Capture-seq, including six mutant isoforms from the two 3′-del *NRXN1*^⁺/⁻^ derived organoids (**Fig. 4f**). In addition, we identified seven β-isoforms, including one mutant β-isoform (**Extended Fig. 4**). Compared with the control line, the mutant α-isoforms were expressed widely across all cell types in both lines with 3′-*NRXN1*^⁺/⁻^ (except for *DLX1/2*-INs in 641). In contrast, the mutant β-isoform was detected in fewer cell clusters, showing consistent expression in excitatory neurons in both 3′-*NRXN1*⁺^/⁻^ lines, and exclusively in TACs in line 581 and RG/AC cells in line 641 (**Fig. 4g**). The composition of *NRXN1* isoform by cell type demonstrated that mutant isoforms from the allele bearing 3’-deletion are expressed in all cell types, contributing to altered synaptic activity in excitatory neurons and *DLX1/2*-INs.

With *NRXN1* splicing profiles available across multiple developmental stages, we next asked which stage the control hCOs most closely resemble in terms of *NRXN1* splicing– the fetal neocortex or the adult PFC. We therefore compared the exon inclusion level across splicing sites (SS1-6) for *NRXN1* in three major cell classes that appear in all samples, *i.e.,* GABAergic inhibitory neurons, excitatory neurons, and RG/AC (**Fig. 4h**). In GABAergic neurons, exon inclusion levels were highly correlated between organoids and the fetal neocortex (Pearson’s *r = 0.97, p = 3.82E-04*) and between cortical organoids and the adult PFC (Pearson’s *r = 0.88, p = 0.008*), although exons at SS2, SS3, and SS5 displayed differential splicing. Similarly, significant agreement was observed between cortical organoids and the fetal neocortex in glutamatergic excitatory (Glu-EX; *Pearson’s r = 0.73, p = 0.04*) and radial glia/astrocyte (RG/AC; Pearson’s *r = 0.78, p = 0.04*) populations, and between organoids and the adult PFC in Glu-EX (Pearson’s *r = 0.88, p = 0.004*) and RG/AC (Pearson’s *r = 0.92, p = 0.003*) populations (**Extended Fig. 4**). Together, these findings indicate that cortical organoid splicing profiles partially recapitulate both developmental and mature splicing patterns, with a subset of splice sites showing cell type – and stage-specific divergence (**Fig. 4h, Extended Fig. 4**).

### Mutant *NRXN1* isoforms are enriched in MLIs in ASD cerebellum with *exonic* deletion

*NRXN1* deletions are also strong risk factors for autism spectrum disorder (ASD)^45,46^. The cerebellum shows consistent structural and functional alterations in ASD, and *NRXN1α* isoform complexity is significantly different in the cerebellum compared to the cerebral cortex. To understand *NRXN1* isoform diversity in human brain with *NRXN1* heterozygous deletion, we applied our probe-based long-read Capture-seq to the postmortem cerebellum of an ASD patient with *NRXN1*^+/-^ (encompassing exon 9-13) (**Extended Fig. 5a**). We performed single-nucleus RNA-seq on this sample and obtained 9,689 high-quality nuclei. Unsupervised clustering and annotation identified eight cerebellar cell type classes: excitatory granule cells; inhibitory neurons (Purkinje cells; two types of molecular layer interneurons MLI1, MLI2; and Purkinje-layer interneurons); astrocytes; Bergmann glia; oligodendrocytes; and OPCs (**Fig. 5a**). These annotations matched expected marker gene expression^47^ (**Extended Fig. 5b, Supplementary Table 5**).

**Figure 5.**
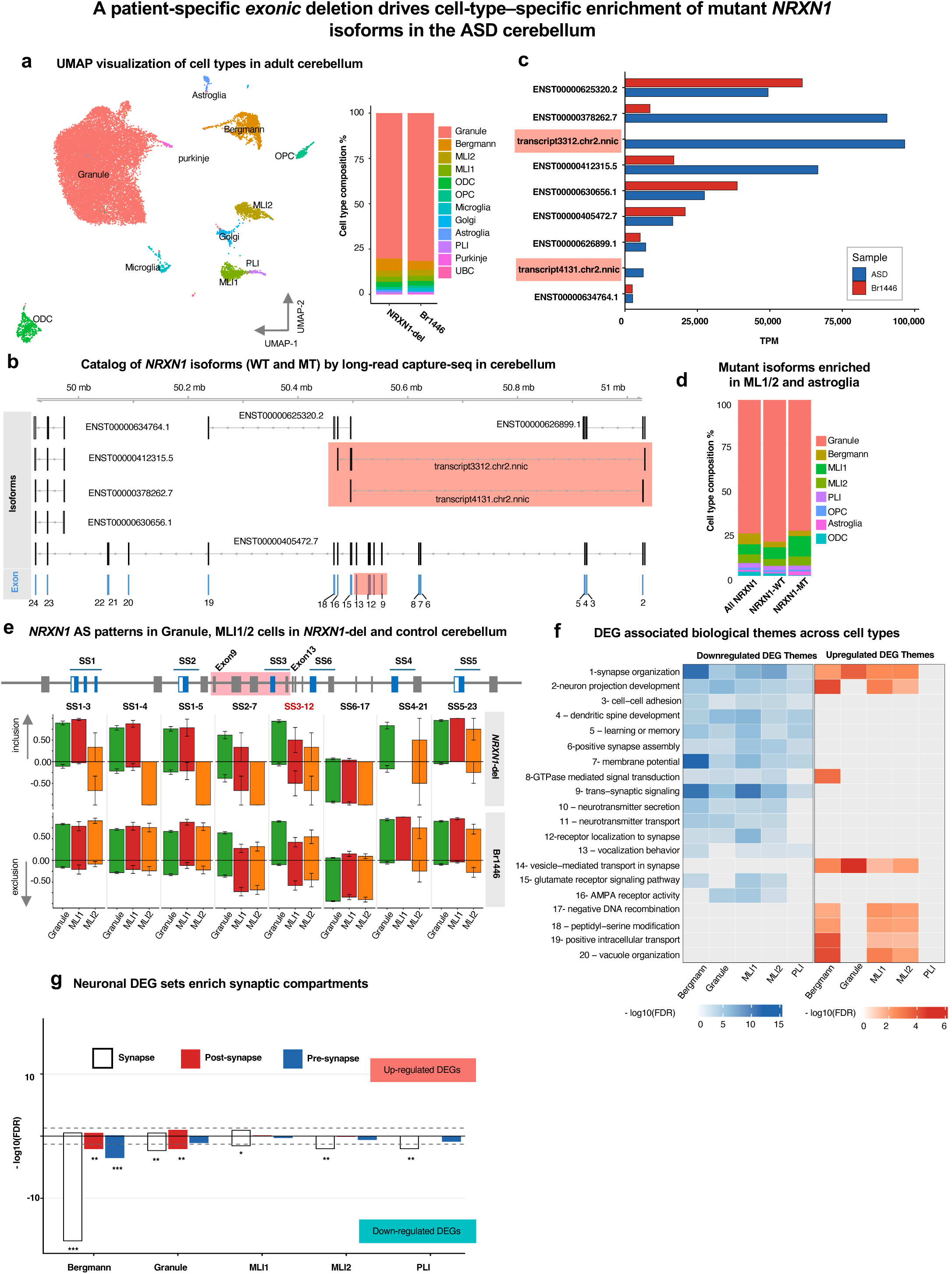
A patient-specific *exonic* deletion drives cell-type–specific enrichment of mutant *NRXN1* isoforms in the ASD cerebellum. (a) *Left:* UMAP of putative cell types in the cerebellum, with colored clusters representing neuronal and non-neuronal subtypes. *Right:* Cell proportion of each cell class in the cerebellum of ASD patient (*NRXN1*-del) and the control. (b) Schematic of *NRXN1* isoform structures identified by long-read capture-seq, with each row representing a unique *NRXN1* isoform. The *NRXN1*^⁺/⁻^ specific mutant isoforms are highlighted in red. (c) Expression level of *NRXN1* isoforms (expression level as TPM) from cerebellum of control or *NRXN1*-del. The red box highlights mutant isoforms specific to the deletion-carrying allele. (d) The proportion of cells in each cell class expressing any *NRXN1* isoforms, wild-type isoforms(*NRXN1*-WT) and mutant isoforms (*NRXN1*-MT). (e) Normalized *NRXN1* splicing events at SS1–SS6 for Granule and MLI1/2 cells from *NRXN1*^+/-^(*NRXN1*-del) and control (Br1446) cerebellum. Schematic splice graph of *NRXN1* depicting splice sites (SS1–SS6), with red shading marking regions impacted by the exonic deletion allele. Alternatively spliced exons at each splice site are highlighted in blue. Exon inclusion is shown upward (“inclusion”) and exclusion downward (“exclusion”). Data represent mean ± SEM. (f) Aggregate enrichment of GO terms within each biological theme across cell types by down-regulated (blue) and up-regulated (red) DEGs, excluding themes implicated by only one term and nonsignificant cell types. (g) Overrepresentation of synaptic compartment genes [SynGO database ^65^] within cell type–specific DEGs. *\*P* < 0.05, *\*\*P* < 0.01, *\*\*\*P* <0.001, Hypergeometric test, FDR < 0.05.

When applying the probe-based long-read Capture-seq to the single nuclei cDNA of the cerebellum, we identified 9 *NRXN1* isoforms with cell barcodes matched to short-read data, including two mutant isoforms associated with the exon 9-13 deletion (**Fig. 5b,c)**. Among *NRXN1* reads with cell barcodes matched to short-read data, the mutant isoforms 3312 and 4131 were exclusively detected in *NRXN1*-deletion cerebellum, with transcript 3312 being the most abundant and 4131 less abundant, together accounting for 28.34% of reads (**Fig. 5c**). When assigning isoforms to cell types, we found the two mutant isoforms were significantly enriched in MLI1 (Fisher’s Exact test, *p* =5.72E-18), MLI2 (Fisher’s Exact test, *p* =0.04), and astroglia cells (Fisher’s Exact test, *p* =51.49E-13) compared to wild-type *NRXN1* isoforms (**Fig. 5d**). The enrichment of mutant isoforms in MLIs, key GABAergic regulators of Purkinje cell output, links these mutations to ASD-associated E/I imbalance, suggesting impaired inhibitory circuitry as a primary site of cerebellar vulnerability^48–50^.

Next, we examined *NRXN1* splicing across six alternative splice sites in these cell types. In the ASD case with *NRXN1* deletion, more than 70% of granule cells showed exon inclusion at SS1, SS3, SS4, and SS5, with moderate inclusion at SS2 (>50%) and predominant exon exclusion at SS6. The two MLI subtypes exhibited distinct splicing patterns, particularly at SS1 and SS2. In controls, *NRXN1* splicing varied across cell types, most prominently at SS2–SS4. Comparative analysis revealed broadly similar splicing patterns between ASD and control in granule cells and MLI1 cells, with high exon inclusion at SS1, SS3, SS4, and SS5, variable inclusion at SS2, and predominant exclusion at SS6. In contrast, subtype-specific differences were observed in MLI2 cells (SS1 and SS2), PLI cells (SS3), and Purkinje cells (SS4) (**Fig. 5e, Extended Fig. 5c**).

To gain insight into how exonic deletion of *NRXN1* affects the DEG networks in the cerebellum potentially associated with ASD, we performed differential expression (DE) analysis for each cell type between control and *NRXN1*-del (see Materials and Methods for details). we then performed Gene Ontology analysis on the up– and downregulated genes per cell type and grouped significantly enriched terms into 20 biological themes (**Extended Fig. 5d**, **Supplementary Table 6,7**). Biological themes related to synapse organization and neuron projection development were associated with DEG sets within excitatory and inhibitory neuronal populations (**Fig. 5f**). Given the prevalence of pathways related to synaptic structure and function, we further assessed enrichment of synaptic compartment genes using the SynGO database. The top-level SynGO category “synapse” was significantly enriched (FDR < 0.05, Hypergeometric test) across all neuronal cell types (**Fig. 5g**).

### Cell-type–resolved evaluation of ASO inhibition of GOF *NRXN1* isoforms in cortical organoids

Antisense oligonucleotides (ASOs), recently used to treat several neurological diseases^51^, provide a strategy to selectively target disease-associated mutant isoforms by modulating specific RNA splice sites. In our recent work, we tested ASO efficacy in 2D iGLUT neurons with bulk RNA-seq, but this approach lacked cellular complexity and cell-type resolution^19^. We hypothesized that ASO efficacy and impact are cell-type specific, particularly in inhibitory neuronal populations enriched for mutant isoforms. Here, we applied the long-read single cell targeted-seq approach to evaluate ASO in three-dimensional (3D) human cortical organoids in a cell type–resolved manner. We adopted the same ASO design that target the exon 20–24 mutant splice junction and applied ASO treatment (10 μM) to the organoids derived from 641 line (3’-del) for 96 hours (see Methods; Extended Fig. 6a, b). We collected the ASO-treated organoids for single-cell cDNA synthesis, using half of the single-cell cDNA for *NRXN1* isoform profiling with both 5’RACR-seq and long-read Capture-seq, while the other half used for cell type annotation by single-cell RNA-seq. From the NRXN1-targeted long-read sequencing data, we found ASO-Aplus decreased total NRXN1-MT isoforms by approximately 51% in all cells (**Extended Fig. 6b,c**). Differential splicing analysis confirmed a reduction in GOF splicing (**Fig. 6a**, ΔPSI (percent spliced index) = −0.23), increased exon23 inclusion at SS5 (ΔPSI = 0.313) and decreased exon 21 inclusion at SS4 (ΔPSI = −0.154), accompanied by a significant shift from α to β isoforms (**Extended Fig. 6c**, p < 0.001, Fisher’s exact test).

**Figure 6.**
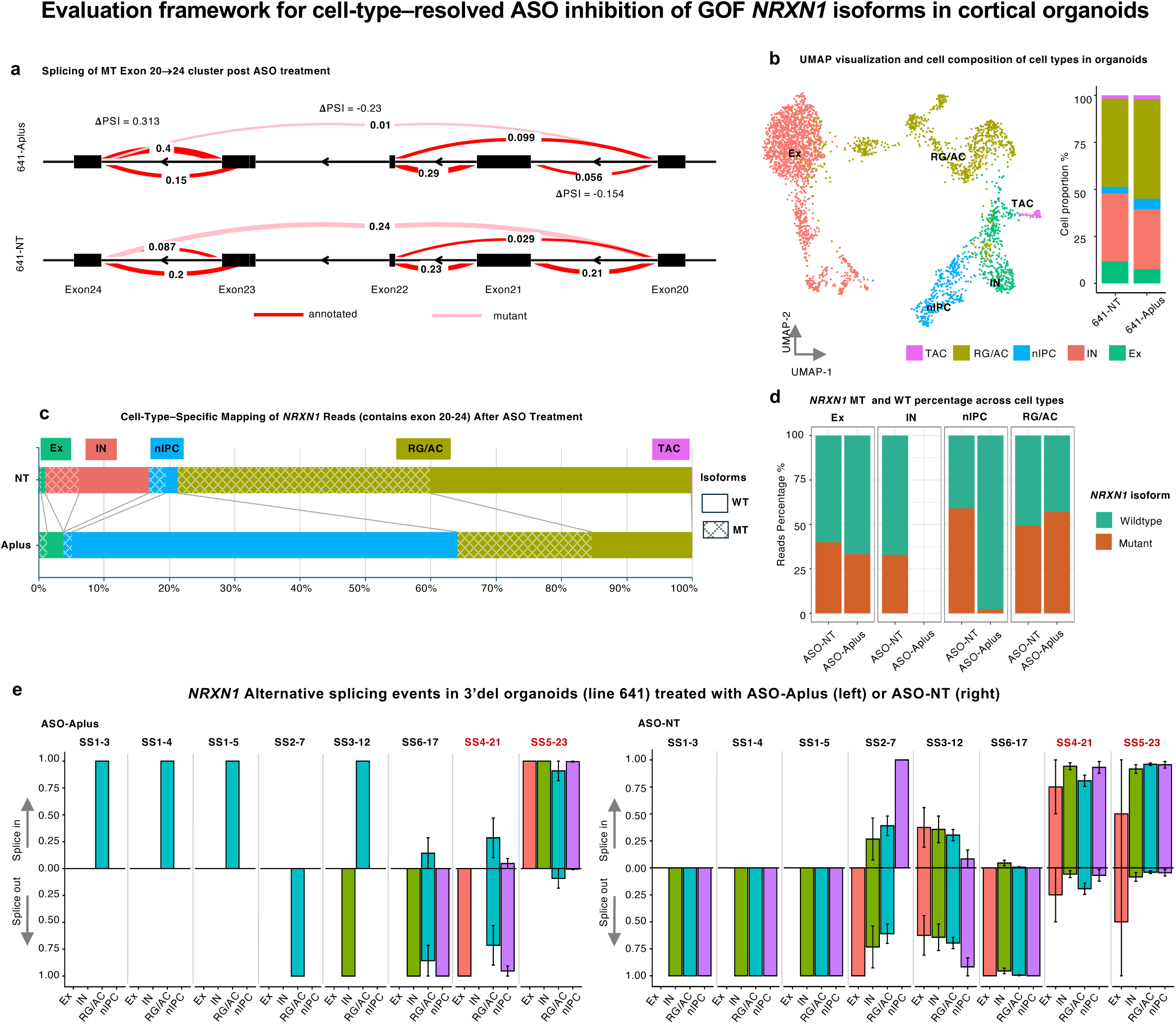
Evaluation framework for cell-type–resolved ASO inhibition of GOF *NRXN1* isoforms in cortical organoids. (a) *Left:* UMAP of putative cell types in ASO-treated hCOs derived from line 641, with colored clusters representing neuronal and glial subtypes. *Right:* Proportions of each cell class in line 641 treated with ASO-NT or ASO-Aplus. (b) Splicegraphs displaying the splicing of the MT exon 20→24 cluster in 641 hCOs treated with ASO-Aplus (upper) or ASO-NT(lower), compared via a Dirichlet-multinomial generalized linear model. (c) Cell-type–specific mapping of NRXN1 reads (containing exons 20–24) following ASO-NT or ASO-Aplus treatment. Color coding of cell types corresponds to the scheme used in the UMAP (Fig. 6a). (d) Percentage of wildtype or mutant *NRXN1* isoforms detected by long-read RACE-seq across cell types in 641 hCOs with ASO treatment. A significant enrichment of the wildtype isoform in nIPCs is observed in the 641-Aplus condition (OR = 0.015, *p* = 1.53 × 10⁻⁷, BH-adjusted *p* = 6.13 × 10⁻⁷, Fisher’s Exact test). (e) Normalized *NRXN1* splicing events at SS1–SS6 for neuronal (EXs and INs) and non-neuronal (RG/AC, nIPC) cells in 641 hCOs treated by ASO-Aplus treated (left) or ASO-NT (right). Exon inclusion is shown upward (“splice-in”) and exclusion downward (“splice-out”). Data represent mean ± SEM.

To understand which cell types are mostly perturbed by ASO targeting of the mutant isoforms in the hCOs, we first analyzed single-cell data from the ASO-treated organoids for cell type annotation (**Fig. 6b**, **Extended Fig. 6d**). We did not observe significant difference for cell composition between the organoids treated with ASO-NT and ASO-Aplus (**Fig. 6b**). Single-cell RNA-seq profiling revealed robust downregulated DEGs in IN and RG/AC cells after ASO-Aplus treatment, enriched for regulation of translation and RNA splicing reported by previous study^52^ (**Extended Fig. 6e,f** and **Supplementary Table 8**).

Integration with cell type–resolved single-cell RNA-seq data revealed that the proportion of *NRXN1* reads containing exons 20–24 was markedly reduced in RG/AC cells and IN cells following ASO-Aplus treatment (**Fig. 6c**). Most importantly, ASO-Aplus significantly reduced proportion of *NRXN1*-MT reads in nIPCs (**Fig. 6d**; OR = 0.015, *p* = 1.53 × 10⁻⁷, BH-adjusted *p* = 6.13 × 10⁻⁷, Fisher’s Exact test) and showed decreased trend in excitatory neurons, not in RG/AC, indicating cell-type–specific differences in the uptake efficiency of ASOs delivered via gymnosis and/or their knockdown efficacy within each cell type. The activity of ASO-Aplus is consistent with established cell-type differences in ASO efficacy in the central nervous system, where neuronal populations show greater target engagement and higher transfection^53,54^. The decreased proportion of RG/AC in ASO-treated conditions without accompanying *NRXN1*-MT knockdown in those cells suggests a selective cytotoxic or antiproliferative effect on RG/AC rather than on-target silencing^55^. Splicing analysis revealed that exon 21 (SS4) was reversely spliced across all cell types, whereas exon 23 (SS5) remained consistently included following ASO-Aplus compared to ASO-NT treatment. In RG/AC cells, exons at SS1, SS3, and SS4 also exhibited reversed splicing, while splicing patterns at SS2, SS5, and SS6 were largely unchanged between ASO-Aplus and ASO-NT conditions (**Fig. 6e**). Notably, neither wild-type transcripts containing exons 20–24 nor the mutant were detected in INs (**Fig. 6c**). This likely reflects partial overlap of the ASO-Aplus sequence with exons 20 and 24, resulting in unintended targeting of wild-type *NRXN1* in addition to *NRXN1*-MT derived from the 3′ deletion allele (**Extended Fig. 6b**).

Altogether, we characterized alterations in MT vs WT *NRXN1* transcripts and splicing events across cell types following ASO treatment using long-read Capture-seq and long-read RACE-seq in hCOs derived from a 3′-del patient. These results highlight the combined use of long-read Capture-seq and RACE-seq as a powerful toolkit for comprehensive evaluation of ASO-induced changes in transcript repertoires and splicing patterns across diverse cell types.

## DISCUSSION

We applied an integrative isoform sequencing strategy combining long-read Capture-seq and long-read RACE-seq to generate a cell-type–resolved catalog of *NRXN1* isoform diversity across human cortical organoids, prenatal cortex, adult PFC, and cerebellum. *NRXN1* exhibited pronounced cell type– and subtype-specific splicing patterns. In human cortical organoids and adult PFC, *NRXN1* splicing showed clear cell-type specificity, whereas analysis of cerebellar tissue from an ASD case revealed deletion-associated isoforms enriched in molecular-layer interneurons and astroglia. Using this framework, we further demonstrated that ASOs targeting the mutant splice junction promote degradation of mutant isoforms in specific cell types within patient-derived organoids. Together, these results establish a platform for resolving *NRXN1* alternative splicing at cell-type resolution and for evaluating ASO-mediated transcript perturbations across cellular contexts.

Our strategy integrates probe-based enrichment, single-cell barcoding, and long-read sequencing to overcome key challenges in detecting low-abundance, full-length *NRXN1* transcripts. Targeted capture increased on-target recovery by approximately 50-fold compared with untargeted long-read approaches while preserving single-cell resolution. This workflow links full-length isoforms to their cellular identities, enabling quantitative mapping of *NRXN1* splicing across neuronal and glial populations and connecting genetic variation to functional transcriptomic phenotypes.

Glutamatergic neurons, the predominant cortical population, demonstrated highly concordant of *NRXN1* splicing patterns between prenatal neocortex and adult PFC. Likewise, glutamatergic neurons in iPSC-derived cortical organoids recapitulated splicing profiles observed in the prenatal cortex, suggesting that *NRXN1*^+/–^ organoids capture key developmental splicing programs. GABAergic interneurons in the PFC exhibited substantial molecular diversity, reflected in distinct *NRXN1* splicing patterns. Although *PVALB*-expressing and *SST*-expressing interneurons both originate from the MGE (*LHX6*+), they showed subtype-specific exon usage consistent with their known functional divergence^56,57^. Notably, *NRXN1* splicing programs appear to be established early in development and remain largely stable, as evidenced by the strong concordance of exon inclusion levels in *PROX1^+^*interneurons between the prenatal neocortex and adult PFC^20^.

Analysis of an *NRXN1*^+/–^ cerebellum from an ASD patient revealed unique mutant isoforms carrying an exon 9–13 deletion that were enriched in molecular-layer interneurons and astroglia, whereas granule cells predominantly expressed wild-type isoforms. Compared to a neurotypical cerebellum, *NRXN1* intergenic deletion was associated with cell-type–specific alterations in gene expression networks. Notably, pathways related to synapse organization and neuron projection development were enriched in both excitatory and inhibitory neuronal populations. Consistent with this, SynGO analysis showed significant enrichment of synaptic compartment genes across all neuronal cell types, indicating convergent disruptions in synaptic structure and function. ^13,59^ These findings support previous reports that *NRXN1* deletions perturb cortical and cerebellar circuits through cell-type–specific mechanisms, consistent with circuit dysfunction observed in ASD and schizophrenia^58–60^. Given that *NRXN1* isoforms determine synaptic partner specificity, the isoform atlas generated here provides a foundation for isoform-targeted therapeutic strategies, including ASO-based modulation of pathogenic splicing.

Long-read Capture-seq and RACE-seq provide complementary approaches for transcript isoform characterization. Capture-seq uses probe-based enrichment of single-cell barcoded cDNA^61^ to detect diverse isoforms, including alternative transcription start and termination sites, across multiple cell types while preserving relative transcript abundance. However, this method preferentially captures abundant transcripts and shows reduced sensitivity for low-abundance or long α-isoforms. In contrast, RACE-seq selectively amplifies transcripts from defined 5′ or 3′ ends, enabling accurate reconstruction of full-length isoforms, including those spanning deletion junctions. Although RACE-seq is less quantitative and has lower throughput, it provides higher sensitivity for rare transcripts. Applying this framework to assess ASO-induced transcript perturbations in organoids yielded fewer than 1,000 cells following treatment with either ASO-Aplus (827 cells) or control ASO-NC (668 cells) (**Supplementary Table 2**). The limited recovery likely reflects ASO-associated toxicity (**Extended Fig. 6a,b**) and reduced cell viability in day-222 organoids^62,63^. Long-read Capture-seq profiling of these samples provided high-resolution transcriptome-wide splicing information; however, sequencing depth was insufficient to comprehensively analyze splicing at sites SS4 and SS5 due to the limited number of recovered cells. In this context, 5′ RACE-seq complemented Capture-seq by enabling detection of mutant isoforms derived from the deletion allele when expression levels were low.

Some limitations should be noted. Both Capture-seq and RACE-seq remain constrained by RNA integrity and read length, particularly in postmortem tissue. In addition, scRNA-seq requires tissue dissociation, which disrupts cell–cell interactions and introduces dissociation bias, limiting the ability to assess how *NRXN1* isoform perturbations reshape local cellular networks in situ. Future integration with spatial transcriptomics could help preserve tissue architecture and resolve cell–cell interactions. Another limitation of this study is the small number of patient-derived samples. Specifically, we analyzed two organoids’ lines carrying 3′-*NRXN1* deletions, along with one ASD cerebellum sample harboring an *NRXN1* deletion and one control. This limited sample size reflects both the rarity and non-recurrent nature of *NRXN1* exonic deletions in the currently available biobank^3,64^. However, building on this study, we are in collaboration with a broad research network of NRXN1 deletion and will expand the research to additional patient-derived samples in future studies.

With the promise of ASO based gene therapy targeting RNA splicing, resolving transcript diversity at cell-type resolution is essential for understanding the delivery and knockdown efficiency. Our targeted long-read sequencing approach enables high-resolution characterization of isoform usage and splicing shifts across distinct cell populations following ASO treatment, providing a cell-type–specific readout of isoform knockdown efficiency. Overall, this study demonstrates that targeted long-read sequencing offers a framework for comprehensive characterization of low-abundance disease-associated transcript isoforms with cell type specificity, with broad application to study natural isoform variations and the evaluation and optimization of ASO-based interventions targeting pathogenic splicing events.

## METHODS

### Data Availability

Raw long-read sequencing data have been deposited in the Sequence Read Archive under accession number PRJNA1447486. Processed and raw single-cell RNA sequencing (scRNA-seq) data, including gene–cell count matrices and associated metadata, have been deposited in the Gene Expression Omnibus under accession number GSE327718 (to be updated upon release).

#### Acquisition and processing of human post-mortem samples

##### The MSSM cohort single nuclei cDNA

The sequencing data and full-length single-nucleus cDNA libraries of 61 individuals from the Mount Sinai School of Medicine cohort were generated from the previous publication^27^. The leftover barcoded full length single-nucleus cDNA was used as input for the current study. To obtain 1-1.5 µg of cDNA for the hybridization step, 5-10 ng of the surplus barcoded 10x cDNA was amplified in two 50 µl reactions, which consist of 25 µL 2X PrimeSTAR GXL premix (Takara, Cat# R051A), 1 µL partial read 1 primer (10 µM, 5’-CTACACGACGCTCTTCCGATCT-3’), 1 µL partial TSO primer (10 µM, 5’-AAGCAGTGGTATCAACGCAGAG-3’). The amplification program is set as follows: 98 °C for 30 seconds, 8 cycles of 98 °C for 10 seconds, 65 °C for 15 seconds and 68 °C for 8 minutes, 1 cycle of 68 °C for 10 minutes. The concentration of PCR products was quantified by Qubit, and additional cycles were added if the yield was lower than 500 ng per reaction. The PCR products were cleaned up with 0.6× AMPure XP beads.

#### *NRXN1^+/-^* cerebellum of human post-mortem brain

Postmortem human cerebellum of the ASD case with a heterozygous *NRXN1^+/-^* mutation were collected and consented through the Department of Pathology at Western Michigan University Homer Stryker MD School of Medicine under WCG IRB protocol #20111080 and the other samples were consented through the National Institute of Mental Health Intramural Research Program under NIH protocol #90-M-0142, and was acquired by LIBD via material transfer agreement.

##### Fetal brain

Two human gestational cortical tissue samples (24 gw and 33 gw) were obtained from previous study ^36^ in accordance with the policies and regulations at the Icahn School of Medicine at Mount Sinai and its Institutional Review Board.

##### Single cell/nucleus full-length barcode cDNA

60mg of the frozen post-mortem tissues were submitted to Yale Center for Genome Analysis for single nuclei isolation. Briefly, single nuclei were automatically extracted from the frozen tissue using the Singulator system, which includes washing nuclei suspension in buffer supplemented with sucrose, and nuclei FACS sorting. The isolated nuclei were then processed using the 10x Genomics 5’GEX V2 to create single nuclei cDNA and matched short-read RNA-seq libraries per manufacturer’s instructions.

#### Patient-derived organoids

Human induced pluripotent stem cell (hiPSC) lines used were validated using methods as previously described. Genome-wide SNP genotyping was performed using Illumina genome-wide GSAMD-24v2-0 SNP microarray at the Children’s Hospital of Philadelphia. Cultures were tested for Mycoplasma contamination and maintained Mycoplasma-free. A total of 3 hiPS cell lines derived from fibroblasts collected from 1 healthy subject (690) and two 3’-del lines from two patients (581 & 641) were used for experiments. Approval for this study was obtained from the Stanford IRB panel, and informed consent was obtained from all subjects.

##### Stem cell maintenance

Human iPSCs were obtained as previously described in Flaherty et al., 2019. Under sterile conditions, 6-well plates (Corning 3516) were coated with 60 ug/mL Geltrex matrix (Thermo Fisher Scientific A1413302) diluted in Dulbecco’s Modified Eagle Medium (DMEM, Gibco 11965-092) per well and incubated for at least 30 min at 37 °C. All hiPSC lines were seeded on coated plates and maintained in StemFlex (StemFlexTM Basal Medium (Gibco A3349301) supplemented with StemFlexTM Supplement (Gibco A3349201)) at 37 °C. After 4 days, hiPSC lines were passaged by removing the media washing cells with PBS. Then cells were incubated in 0.5 µM EDTA in PBS (Invitrogen AM9261) for 4 min at 37 °C. After removing EDTA, colonies were broken into smaller pieces by triturating using a p1000 pipette and StemFlex. It is worth noting that very small colonies have a very low survival chance and should be supplemented with 50nM chroman (CHR; MedChemExpress, Cat. No. HY-15392). After dissociation, colonies were distributed in new coated plates containing StemFlex in a ratio of 1:6 (250K cell per well of a 6 well plate). A complete media change was performed the following day. Complete media changes were performed as needed or every other day.

##### Cortical organoid generation

Cultures of hiPSC were dissociated at 70% confluency and checked for signs of differentiation. Mature undifferentiated cells were washed with PBS, followed by incubation with Accutase (STEMCELL™ Technologies, Cat. No. 07920 and CHR for 10-20 min to achieve complete detachment. Cells were gently triturated to achieve a single-cell suspension. The resulting cell suspension was transferred into DMEM (Gibco, Cat. No. 11960044) and centrifuged at 500 × g for 4 min. Cells were resuspended in StemFlex™ medium (Thermo Fisher Scientific) supplemented with CHR and diluted at a density of 1.5 × 10^6 cells/mL. AggreWell™ plates (STEMCELL Technologies) were prepared with anti-adherent solution according to the manufacturer’s instructions. A volume of 2 mL of the hiPSC suspension was seeded per well to promote spheroid formation overnight (defined as day 0). Neural induction was subsequently performed following the human forebrain spheroid protocol described by Sloan *et al*^66^. In brief, medium was replaced daily until day 15, every other day from days 15–42, and every 4 days from day 43 onward. On day 1, embryoid bodies were dislodged and transferred to ultra-low adhesion 10 cm dishes containing Neuronal Induction Media (StemFlex™ medium without StemFlex™ supplement), SMAD inhibitors SB431542 (10 µM; R&D Systems/Tocris Bioscience,1614), LDN193189 (100 nM; STEMCELL Technologies, 1614), XAV (100nM; STEMCELL Technologies,72672). Plates were placed on an orbital shaker at 53 rpm and maintained at the same speed until day 6. From days 6–25, Neuronal Differentiation Media (Neurobasal-A (Thermo Fisher Scientific), 1x Antibiotic-Antimycotic (Gibco,15240062), 1x GlutaMAX™(Gibco, Cat. No. 5050061), and 1x B-27 supplement minus vitamin A (Gibco, Cat. No. 2587010) was used supplemented with recombinant human FGF2/bFGF (20 ng/mL; R&D Systems, Cat. No. 233-FB-01M) and recombinant human EGF (20 ng/mL; R&D Systems, Cat. No. 236-EG). Between days 25–42, Neuronal Differentiation Media was supplemented with brain-derived neurotrophic factor (BDNF; 100 nM; R&D Systems, Cat No.248-BD-025) and neurotrophin-3 (NT-3; 100 nM; PeproTech, Cat No.450-03). From day 42 onwards, organoids were maintained in Neuronal Differentiation Media.

##### Cryoprotection and immunocytochemistry

Organoids were washed with PBS and fixed in 4% paraformaldehyde (PFA)/phosphate buffered saline (PBS) for 3 to 4 hours overnight at 4°C. Organoids were then washed in PBS and transferred to 25% sucrose/H_2_O for 48 hours. Organoids were embedded in gelatin/sucrose embedding solution by dissolving 7.5 g gelatin in 100 ml 10% sucrose solution at 37°C to obtain a 7.5% gelatin/10%sucrose embedding solution. For immunofluorescence staining, 14-16 μm-thick sections were washed with PBS and blocked in 10% donkey serum, and 0.1% Triton-X in PBS for 1 h at room temperature followed by primary (overnight, 4 °C) and secondary antibody (1 h at room temperature, Jackson ImmunoResearch or ThermoFisher Scientific) incubation. Slides were then mounted (VECTASHIELD, Vector Labs) and imaged on a Zeiss microscope with an apotome module equipped with ZEN 3.3 (ZEN pro) software.

Primary antibody list:. Primary Antibody list: *BCL11B/CTIP2* (rat, 1:500, Abcam), *FOXG1* (rabbit, 1:200, Abcam), *GAD1/GAD67* (mouse, 1:1000, Millipore), *HuC/D* (mouse, 1:500, Invitrogen), *KI67* (rabbit, 1:500, Vector Labs), *PAX6* (mouse, 1:200, BD Bioscience), *SOX1* (goat, 1:20, R&D Systems), *TBR1* (rabbit, 1:1000, Abcam).

##### Antisense oligonucleotide treatment

A single HPLC-purified antisense oligonucleotide (ASO) targeting the mutant exon 20/24 splice junction was designed and synthesized by IDT. The ASO contains a phosphorothioate-modified backbone with incorporated locked nucleic acid (LNA) nucleotides (Affinity Plus, Aplus). A non-targeting ASO (ASO-NT) was used as a matched control in all experiments. The sequence of ASO-Aplus as +G*+G*+T*T*G*G*C*T*G*C*A*G*G*+G*+T*+A; and the sequence of ASO-NT as A*A*C*A*C*G*T*C*T*A*T*A*C*G*C. To optimize delivery conditions and dosage, human cortical organoids (hCOs) derived from line 641 (3′-*NRXN1*⁺/⁻) at day 222 were treated with 1 μM or 10 μM ASO-Aplus or ASO-NT, delivered either via Lipofectamine RNAiMAX^67^ or gymnosis, and incubated for 96 h prior to qPCR validation or single-cell and RNA collection.

Expression levels of mutant and wild-type *NRXN1* transcripts were quantified by qPCR to assess ASO knockdown efficiency.

##### Single-cell dissociation and RNA-seq library preparation

Organoids with optimal morphology from each line were selected in days 25, 60, and 120. Five organoids from each line were pooled at each time point and transferred to a 5 cm plate dish and washed with PBS. Organoids were chopped into small pieces using a sharp sterile blade. 300 ul of Accumacs was added on top of the pieces without scattering them. Pieces were collected by a cut tip and were transferred into a 1.5 ml tube. The tubes were incubated at 37°C and 800 g rotation on a thermal shaker for 20 min. After 20 min, cell suspension was very gently pipetted up to 5 times with a p 1000 tip, to facilitate the cell dissociation. 500 ul fresh Accumacs was added into the suspension and incubated again for 20 min. Cell suspension was pipetted up to 5 times more and passed through a filter in a 15 ml sterile tube containing 1 ml of HEPES buffer. The filter was washed with 1 ml of HEPES buffer. The single cell suspension in a 15 ml tube was centrifuged at 300 g for 4 min to collect cells in a pellet. After removing the supernatant, the pellet was resuspended in 200 ul HEPES buffer and cells were counted using a hemocytometer. The suspension was diluted to reach 1200 cell per ml and was submitted to Yale Genomics Core facility for cDNA library prep using 10x 5’ NEXT GEM V2 according to the manufacturer’s instructions.

#### NGS sequencing of single cell/ nucleus cDNA

The prepared 10X Genomics single cell/nucleus libraries were submitted to Yale Center for Genome Analysis and sequenced on an Illumina NovaSeq sequencer with 150-bp paired-end sequencing on an S4 flow cell. For each sample, ∼10,000 cells were sequenced at a minimum depth of 50,000 reads per cell.

#### Probe-panel design

IDT Lockdown probes (Integrated DNA Technologies) are 120-nt long oligos that are biotinylated at their 5’ ends and were designed using the xGen Panel Design Tool (https://www.idtdna.com/pages/tools/xgen-hyb-panel-design-tool) to ensure robust coverage of the targeted regions. Briefly, a customized *fasta* file containing all Ensembl^26^-annotated exons (across all isoforms) of target genes, including UTRs was uploaded as input file. The probe sequence list was generated at the 1x probe tilling density using “Homo sapiens (Human) NCBI GRCh37.p13 (hg19)” as the reference (Supplementary table 1).

The probe panel was further synthesized employing TEQUILA-seq^21^. First, the probe panel pool of 150-nt long oligos, each comprising a 3’ end 30-nt universal primer binding sequence (5’-CGAAGAGCCCTATAGTGAGTCGTATTAGAA-3’) followed by a 120-nucleotide target-specific probe, was synthesized with Twist Bioscience (Supplementary table 1). Then the oligo pools were amplified and biotin-labeled using nickase-induced linear strand displacement amplification as described previously. Finally, synthesized probes were purified with 1.8× volumes of AMPure XP beads and quantified by NanoDrop 2000 Spectrophotometer.

#### Long-read Capture-seq on single cell/ nucleus cDNA

All hybridization and capture steps were performed following the IDT “Hybridization capture of DNA libraries using xGen Lockdown probes and reagents” protocol. Briefly, 1-1.5 µg of amplified 10x cDNA together with 1 µL of each blocking oligos (1 mM, **Supplementary Table1**) were dried using a speed vac (Thermo, Cat# SPD140P1). Next, the dried sample were re-hydrated with 8.5 μL of 2× Hybridization Buffer, 2.7 μl Hybridization Buffer Enhancer (xGen IDT 1080577), and 3.8 μL nuclease-free water, followed by denaturation at 95 °C for 10 min. The denatured mixture was then incubated with 200 ng of TEQUILA probes at 65 °C for 16 h. Next, the capture step was performed by adding 100 μL of washed M-270 streptavidin beads (Invitrogen, #65306) to the mixture, followed by incubation at 65 °C for 45 minutes with vortexing every 10 minutes to ensure thorough mixing. After capture, the mixture was then immediately washed with high-temperature and room temperature buffers and resuspended in 40 μL nuclease-free water.

The streptavidin bead-captured cDNA was amplified using PrimeSTAR GXL premix (Takara, Cat# R051A) and 10x cDNA amplification primers by incubating at 95 °C for 3 min, followed by 12 cycles of (98 °C for 20 s, 65 °C for 15 s, 68 °C for 8 min), with a final extension at 68 °C for 8 min. Post-capture PCR products were purified using 0.7× volumes of AMPure XP beads, and quantified by the Qubit dsDNA High Sensitivity assay and size checked by Agilent TapeStation.

#### Gene-targeted single cell/ nucleus RACE-PCR

To sequence the *NRXN1α* isoforms and 10x single cell barcode from one long-read, the modified RACE-PCR was employed to amplify the target from the 10x Genomics 3’GEX or 5’GEX cDNA. 5’RACE was used for cDNA generated by 10x 5’GEX kit, while 3’RACE was used for cDNA generated by 10x 3’GEX kit.

##### PCR primer design

As the expression level of *NRXN1α* is relatively low in adult brains, nested PCR would be needed to achieve enough molecules. The gene-specific primers (GSPs) should meet the criteria: 23-28 nt long to ensure specific annealing, with 50%-70% GC, a T_m_ 65°C (best to have a T_m_ >70°C) and be specific to the target gene (**Supplementary Table 1**).

Universal primers: The universal primer for first –round PCR should be the partial read1 sequence from 10x cDNA. To increase sensitivity for the second-round PCR, a sequence of “AATGATACGGCGACCACCGAGATCT” was added to the 5’ of the read1 (**Supplementary Table 1**), forming the second-round universal PCR primer (UP2) (**Supplementary Table 1**).

##### Protocol

The input of 1-50 ng of 10x 5’GEX or 3’GEX cDNA was used for the RACE-PCR. The first round of RACE-PCR is carried out with 1 µL partial Read1 primer (10 µM) and 4 µL *NRXN1* –specific primer 1 (10 µM) in a 50 µl-reaction, which consists of additional 25 µL 2X PrimeSTAR GXL premix (Takara, Cat# R051A), 1-50 ng of full-length single cell cDNA by 10x 5’GEX or 3’GEX kit. Choose the PCR program based on the GSP T_m_. Program 1: when GSP T_m_ >70°C, run 5 cycles of 94°C for 30s, 72°C for 8min; 5 cycles of 94°C for 30s, 70°C for 30s and 72°C for 8min; 15-20 cycles of 94°C for 30s, 68°C for 30s and 72°C for 8min. Program 2: when T_m_ = 60-70°C, run 15-20 cycles of 94°C for 30s, 68°C for 30s and 72°C for 8min. The concentration of PCR products was quantified by Qubit, and additional cycles can be added if the concentration of PCR product is lower than 10 ng/µL. After the PCR, 3 µL of products were analyzed with 1% agarose/ GelRed gel (biotium, cat# 41002), while the rest products were purified using 0.7× volumes of AMPure XP beads and eluted with 30 µL nuclease-free water. If the primary PCR fails to show the distinct band(s) of interest or produces a smear, a second-round of RACE-PCR should be performed.

The second-round of RACE-PCR is performed with 2.5 µL of UP2 (10 µM) and 2.5 µL *NRXN1*-specific primer 2 (10 µM) in a 50 µl-reaction together with 25 µL 2X PrimeSTAR GXL premix and 10 µL of the purified first-round RACE-PCR products. The PCR program follows the program 2 and runs for 10-15 cycles based the final concentration of PCR products. After the PCR, 3 µL of products were analyzed with 1% agarose/ GelRed gel for size check, which is followed by the purification using 0.7× volumes of AMPure XP beads.

#### Library preparation for Oxford Nanopore sequencing

Oxford Nanopore SQK-NBD114.24 was used for library preparation. Amplified cDNA from post-capture of RACE-PCR, approximately 200 ng of PCR products from each sample were used for library preparation according to the ONT SQK-NBD114.24 protocol. Briefly, cDNA products were end-repaired and dA-tailed with NEBNext Ultra II End Repair/dA-Tailing Module by incubating at 20 °C for 30 min and 65 °C for 30 min. The cDNA was then purified with 1× volume of AMPure XP beads and eluted in 10 μl of nuclease-free water. Native barcode ligation was then performed using 10 μl of end-prepped DNA, 2.5 μl of native barcodes, and 12.5 μl of Blunt/TA Ligase Master Mix (NEB, #M0367) by incubating at room temperature for 30 min. After the ligation step, 2.5 μl of EDTA was added to each tube to inactivate the ligase. Barcoded cDNA samples were then pooled and purified using 0.4× volume of AMPure XP beads and eluted in 31 μl of nuclease-free water. Next, ligation of Native Adapter was performed with 30 μl of pooled barcoded samples, 5 μl of native adapter, 10 μl of 5× NEBNext Quick ligation reaction buffer, and 5 μl of NEBNext Quick T4 DNA ligase at room temperature for 30 min, followed by the bead’s purification using 0.45× volume of AMPure XP beads and twice wash with short fragment buffer. The libraries were quantified with Qubit, and 50 fmol of final library was sequenced with a R10.4.1 MinION or PromethION flow cell for 72 hours.

#### Single-cell RNA-seq processing and analysis

Raw 10x Genomics single-cell RNA-seq data were processed using Cell Ranger (v7.2.0) with the GRCh38 reference genome to generate gene-cell UMI count matrices. Downstream analyses were performed in Seurat (v5.0.3)^68^ under R (v4.4.0). Cells with fewer than 200 detected genes, more than 6,000 detected genes, or mitochondrial transcript fractions exceeding 20% were excluded. Potential doublets were identified and removed using DoubletFinder^69^. See the sequencing summary in **the Supplementary Table 2**.

Post-quality control, gene expression values were normalized by scaling each cell’s total expression to 10,000 counts and log-transforming the results. Data were then processed using SCTransform^68^, which performs regularized negative binomial regression to normalize and correct for sequencing depth and mitochondrial content. Highly variable genes were selected for PCA, and dimensionality reduction was performed using UMAP.

Clustering was conducted using a shared nearest neighbor (SNN) graph and the Louvain algorithm to identify transcriptionally distinct populations. Cell type annotation was guided by canonical marker gene expression and cross-referenced with public human brain (or tissue-specific) single-cell catalogs. Differential expression analyses between groups or clusters were performed using the Wilcoxon rank-sum test with Benjamini-Hochberg correction for multiple testing (FDR < 0.05).

Following dimensionality reduction and clustering, known marker genes were used to guide the assignment of biological cell identities. Each cluster was annotated based on the expression of established cell-type-specific markers reported in previous studies.

GO Biological Process (BP) enrichment analysis was performed for each cell type using differentially expressed genes (DEGs) identified between ASD and control samples. Up– and downregulated DEGs were analyzed separately. Enrichment was tested using clusterProfiler (v4.12.6) with a background of all genes tested in that cell type (18,903 genes). Significant GO terms were retained at FDR < 0.05 (Benjamini–Hochberg correction). For each cell type and direction, up to the top 200 most significant terms were carried forward for downstream clustering. To reduce redundancy among significant GO BP terms and identify broad biological themes, semantic similarity clustering was performed using the rrvgo R package. For each direction (up/down), a pairwise semantic similarity matrix was computed across all significant GO terms pooled from all cell types. Terms were clustered into representative parent terms using reduceSimMatrix with a similarity threshold of 0.7. Each GO term was scored as the maximum −log₁₀(FDR) observed across all cell types. The resulting clusters were summarised into biological themes, and the top 20 themes ranked by the number of cell types with significant signal are displayed. For each cell type × theme combination, the cell in the heatmap represents the maximum −log₁₀(FDR) across all member GO terms within that cluster. Enrichment of DEGs within synaptic compartments was assessed using the SynGO database (v1.3). Three cellular component categories were tested: Synapse, Presynapse, and Postsynapse. For each cell type and direction (up/down), a one-sided hypergeometric test. Multiple testing correction was applied using the Benjamini–Hochberg method across all cell type × compartment combinations within each direction. Enrichments with FDR < 0.05 were considered significant.

#### Long-read isoform sequencing analysis

Full-length transcript isoforms were characterized using long-read RNA sequencing (Oxford Nanopore or PacBio). Raw reads were basecalled using dorado (v0.4.1). Reads were quality-filtered to retain those with a minimum mean Phred quality score of Q10 and minimum read length of 200 bp, and adapter sequences were trimmed prior to alignment. Processed reads were aligned to the human reference genome (GRCh38) using minimap2 (v2.28) with cDNA splice – aware parameters (-ax splice –-secondary=no –C5). Alignments were coordinate-sorted and indexed using SAMtools (v1.21), and only primary alignments with a minimum mapping quality of MAPQ ≥ 10 were retained for downstream analysis. Isoform reconstruction and quantification were performed using IsoQuant (v3.2) with GENCODE v43 as reference annotation. IsoQuant^70^ clusters reads into high-confidence transcript models, classifying each as known, novel in catalog (NIC), or novel not in catalog (NNC) based on splice junction composition. Only isoforms supported by at least three full-length reads were retained.

Differential splicing analysis was performed using LeafCutter^71^. Briefly, splice junctions were extracted from BAM files using RegTools, with a minimum anchor length of 8 bp and intron size ranging from 50 to 500,000 bp. Junction files from all samples were combined and clustered into intron groups using the LeafCutter clustering pipeline, requiring a minimum of 50 supporting reads per cluster.

#### Integration of long-read isoform data with short-read single-cell transcriptomes

Integration between long-read and short-read data was performed at both the gene and isoform levels. Cell barcode and UMI assignment from Nanopore reads was accomplished using BLAZE (v1.4.0)^72^, which accurately detects 10x cellular barcodes and unique molecular identifiers (UMIs) directly from Nanopore cDNA reads. BLAZE localizes barcode sequences within read adapter contexts, compares candidate sequences against the 10x whitelist, and assigns valid barcodes with up to one mismatch tolerance to each read. This enabled unambiguous association of long reads with individual cell identities and facilitated construction of a cell × isoform count matrix for isoform-level single-cell analysis.

### *NRXN1* Isoform Quantification using *miniQuant*

For the BA4 samples, *miniQuant*^73^ was applied in hybrid mode, integrating both short-read and long-read RNA-seq generated in-house to improve isoform-level quantification. For the FC and hippocampus datasets^25,74^, where only long-read RNA-seq data were available, *miniQuant* was applied in long-read-only mode to estimate isoform abundances.

In *miniQuant*, long-read cDNA sequencing provides high-confidence isoform structures and reduces ambiguity in assigning reads to specific isoforms, whereas short-read sequencing improves quantification accuracy for low-abundance transcripts by reducing sampling error. *miniQuant*’s gene-specific weighting model adaptively balances these complementary data sources based on isoform structural complexity (K-value), transcript abundance, and sequencing depth, producing optimized isoform abundance estimates. Isoform expression levels were reported as transcripts per million (TPM) to allow direct comparison across samples.

To ensure quantification reliability, we filtered out low-confidence isoforms following the criteria described in the original *miniQuant* framework, requiring both sufficient long-read support and a minimum expression threshold. Only isoforms with TPM ≥ 1 in at least one sample were retained for downstream analyses.

### Data visualization and statistical analysis

All visualizations, including UMAPs, dot plots, violin plots, and bar plots, were generated in *R* or *Python* using the *ggplot2*, *matplotlib*, and *seaborn* libraries. For continuous variables, such as comparisons of AS event inclusion levels or isoform expression differences, we applied *Welch’s t-test*, which does not assume equal variances between groups.

## Supporting information

Supplementary Figures with captions

## ACKNOWLEDGEMENTS

We thank Daniel Weinberger and the Lieber Institute for Brain Development at Johns Hopkins School of Medicine, for sharing post-mortem brain tissues. We thank Feng Wang and Yi Xing (Children’s Hospital of Philadelphia) for technical assistance with TEQUILA-seq. This work is supported by the National Institute of Mental Health grants RO1 MH125579 (GF and KJB) and RO1 MH121074 (KJB and GF), and U01MH116442 (PR) and R01MH125246 (PR). Research reported in this publication was supported by the National Institute of General Medical Sciences of the National Institutes of Health under Award Number 1S10OD030363-01A1. N.A.L.H. was supported by the National Institute for Health and Care Research (NIHR) Oxford Health Biomedical Research Centre. This work was also supported in part through the computational resources and staff expertise provided by the Department of Scientific Computing at the Icahn School of Medicine at Mount Sinai.

## AUTHOR CONTRIBUTIONS

LC, YF and GF designed the methods. LC and YF performed targeted long-read sequencing and conducted data analysis. SG generated cortical organoids, and JM performed ASO treatments. YZ assisted with bioinformatic analyses. MBF designed the ASO. JB, JF, and PR provided single-nuclei RNA sequencing data from adult post-mortem brains and contributed to data analysis. SIR and NT provided post-mortem fetal brain tissues. NH provided technical assistance with Capture-seq. GD, KGB, and RS assisted with long-read sequencing. BZ and SM provided technical assistance with single-nuclei cDNA preparation. EAM contributed to data analysis and interpretation. LC, YF, SG, KJB and GF wrote the manuscript with input from all authors. KJB and GF supervised the research.

