## Supplementary Figures with captions for "Cell-type-resolved *NRXN1* isoforms in human brain and hiPSC cortical organoids"

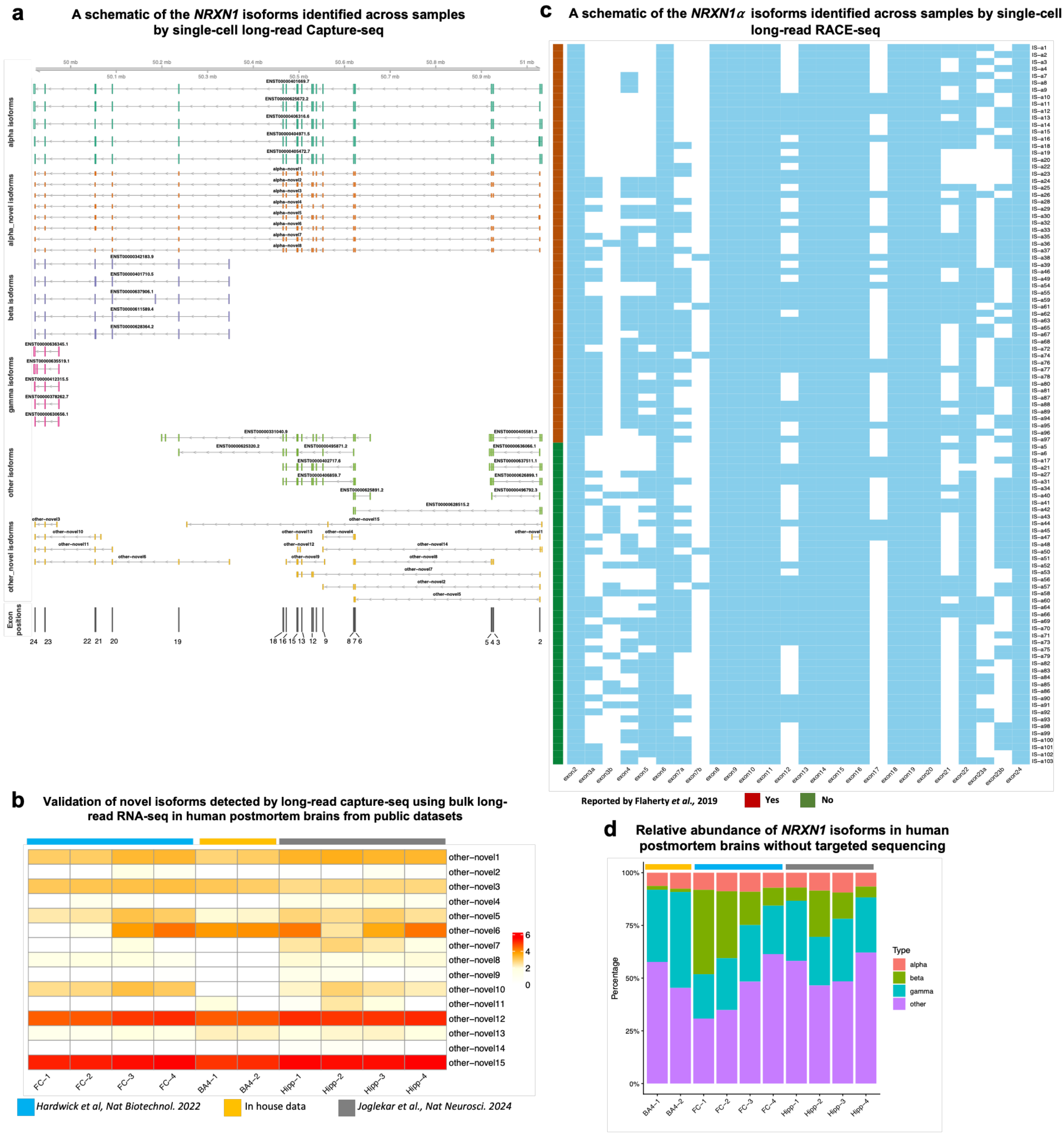

**Extended Figure 1. *NRXN1* isoforms detected by targeted and non-targeted long-read strategies.** (a) Schematic representation of *NRXN1* isoforms identified across samples by single-cell long-read Capture-seq. Each row represents a unique isoform, color-coded as α, α-mutant, β, γ, known others (other), and novel isoforms (other-novel). (b)Expression heatmap of novel *NRXN1* isoforms identified by long-read Capture-seq or bulk long-read RNA-seq in human postmortem brains from public datasets and in-house data; “other” isoforms--transcripts that do not correspond to canonical α, β, or γ isoforms. Expression of each isoform is quantified by *miniQuant*[28] and shown as $\log_{10}$TPM (Transcripts Per Million). (c) Schematic representation of *NRXNα 1* isoforms identified across samples by single-cell long-read RACE-seq. Each row represents a unique *NRXN1α* isoform, with blue exons indicating inclusion and blanks indicating exclusion. Red blocks denote isoforms previously reported, and green blocks denote isoforms identified in this study. (d) The proportion of each *NRXN1* isoform in human postmortem brains from public datasets[25,27] and in-house data.

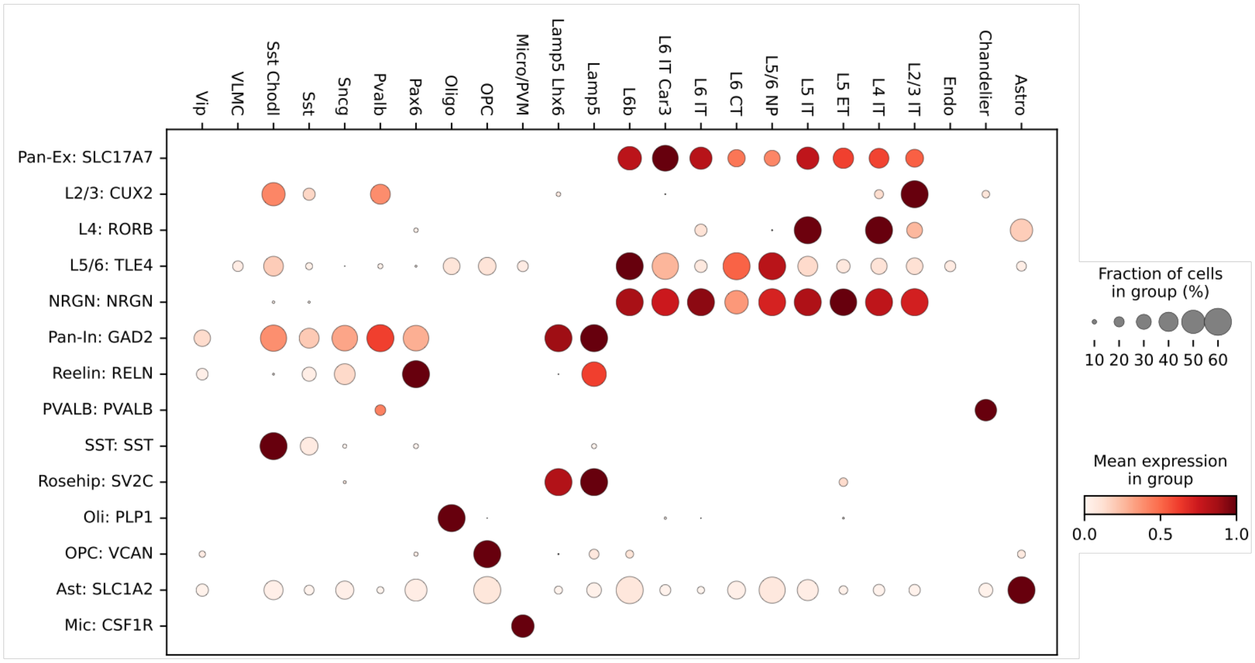

**Extended Figure 2. Dot plot of marker gene expression used for cell-type annotation.** Each column represents a marker gene and each row represents a cell class; darker red indicates higher expression.

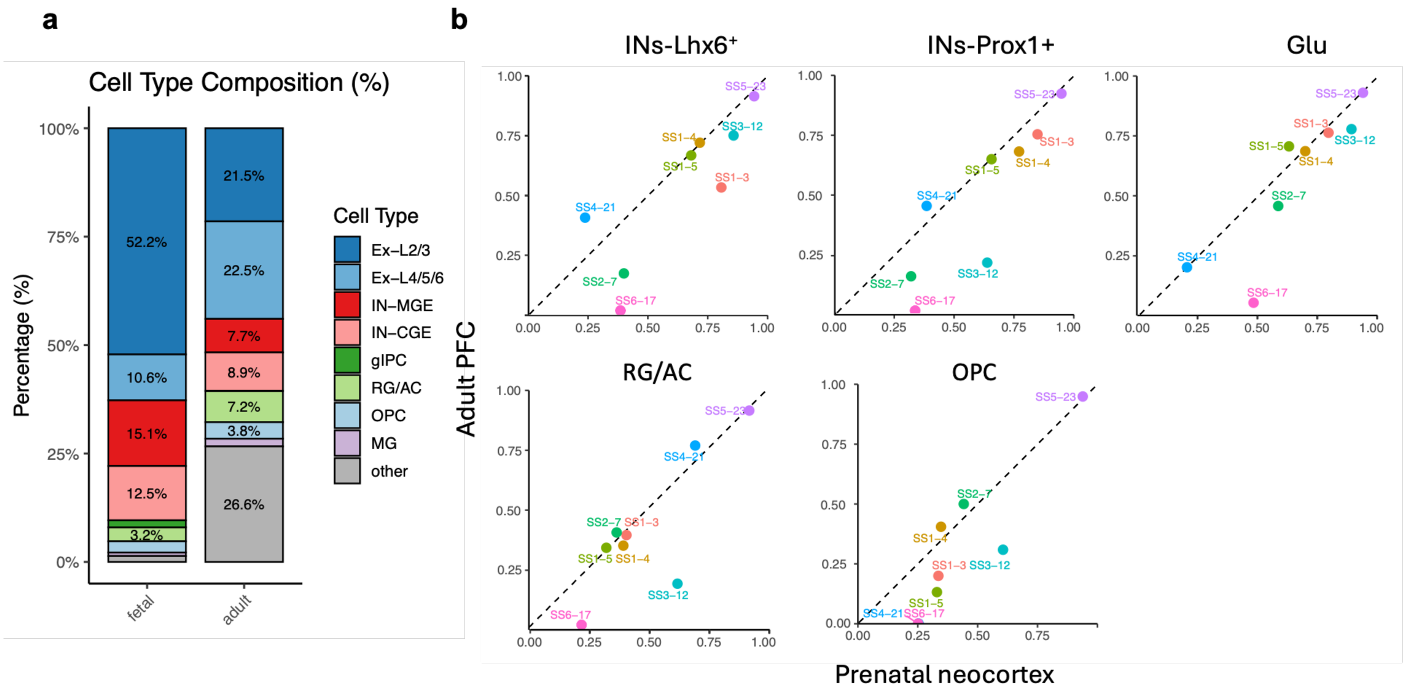

**Extended Figure 3. Cell composition and AS patterns differences between adult PFC and fetal neocortex.** (a) The cell type composition (%) of fetal prenatal and adult PFC. (b) *NRXN1* exon inclusion level across 6 splicing sites between prenatal neocortex and adult PFC in matched cell types. A-axis, *NRXN1* exon inclusion level in prenatal neocortex; y-axis, *NRXN1* exon inclusion level in adult PFC.

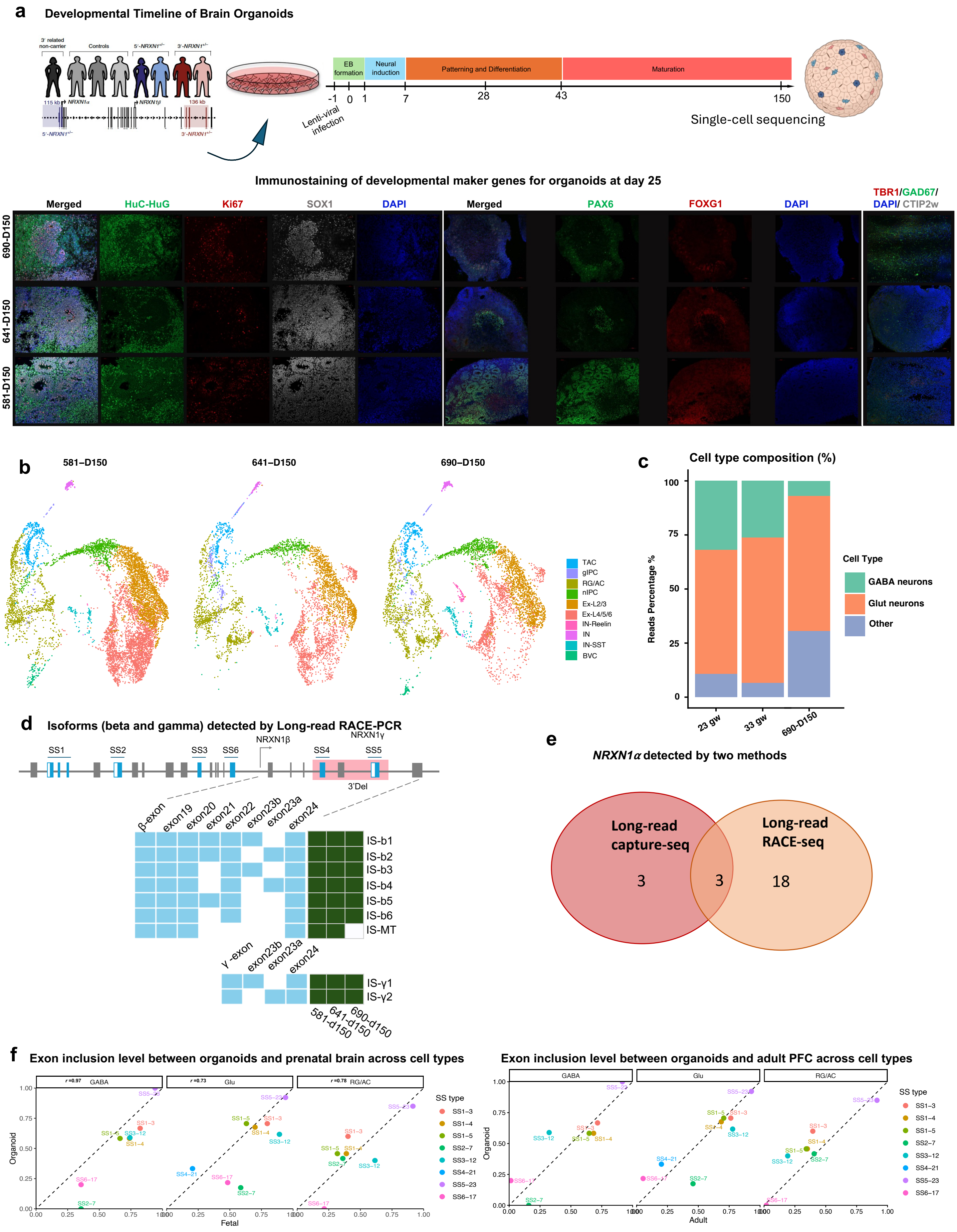
a
Developmental Timeline of Brain Organoids
3' related non-carrier
Controls
5'-NRXN1+/-
3'-NRXN1+/-
115 kb
NRXN1α
NRXN1β
136 kb
5'-NRXN1+/-
3'-NRXN1+/-
EB formation
Neural induction
Patterning and Differentiation
Maturation
-1
0
1
7
28
43
150
Lenti-viral infection
Single-cell sequencing
Immunostaining of developmental maker genes for organoids at day 25
Merged
HuC-HuG
Ki67
SOX1
DAPI
Merged
PAX6
FOXG1
DAPI
TBR1/GAD67/DAPI/CTIP2w
690-D150
641-D150
581-D150
b
581–D150
641–D150
690–D150
TAC
gIPC
RG/AC
nIPC
Ex-L2/3
Ex-L4/5/6
IN-Reelin
IN
IN-SST
BVC
c
Cell type composition (%)
Reads Percentage %
100
75
50
25
0
23 gw
33 gw
690-D150
Cell Type
GABA neurons
Glut neurons
Other
d
Isoforms (beta and gamma) detected by Long-read RACE-PCR
SS1
SS2
SS3
SS6
NRXN1β
SS4
NRXN1γ
SS5
3'Del
β-exon
exon19
exon20
exon21
exon22
exon23b
exon23a
exon24
IS-b1
IS-b2
IS-b3
IS-b4
IS-b5
IS-b6
IS-MT
γ-exon
exon23b
exon23a
exon24
IS-γ1
IS-γ2
581-d150
641-d150
690-d150
e
NRXN1α detected by two methods
Long-read capture-seq
Long-read RACE-seq
3
3
18
f
Exon inclusion level between organoids and prenatal brain across cell types
r=0.97 GABA
r=0.73 Glu
r=0.78 RG/AC
Organoid
Fetal
SS type
SS1−3
SS1−4
SS1−5
SS2−7
SS3−12
SS4−21
SS5−23
SS6−17
Exon inclusion level between organoids and adult PFC across cell types
GABA
Glu
RG/AC
Organoid
Adult
0.00
0.25
0.50
0.75
1.00

**Extended Figure 4. human organoid generation and characterization.**

(a) *Upper*: developmental timeline of human cortical organoids. *Lower*: Immunostaining in hCOs derived from control (line 690) and two 3'-del *NRXN1* hCOs for the developmental marker genes at day 25. (b) UMAP visualization of putative cell types in control hCOs (line 690) and 3'-del *NRXN1* hCOs (line 581 and 641; n=5 organoid per batch, 1 differentiation ) at day 150. (c) Bar graph showing the proportions of each cell class in hCOs at day 150. (d) *Left:* Schematic of *NRXN1*$\beta$ and *NRXN1*$\gamma$ isoform structures identified by long-read RACE-seq, along with a schematic of *NRXN1* gene structure. Each row represents a unique *NRXN1*$\beta$ or *NRXN1*$\gamma$ isoform, with blue blocks indicating exon inclusion and blank spaces indicating exon exclusion. *Right:* Presence or absence of each *NRXN1*$\beta$ (labeled as "IS-b1", *etc*.; MT, mutant *NRXN1*$\beta$ )or *NRXN1*$\gamma$ (labeled as "IS-$\gamma$*1*" and "IS-$\gamma$*2*"),  isoform across 3'-deletion and control hCOs (green = present; blank = absent). (e) Venn diagram shows the *NRXN1*$\alpha$ isoforms detected by long-read capture-seq and long-read RACE-seq. (f) *Left: Pearson's correlation* of *NRXN1* exon inclusion levels across splicing sites (SS1–SS6) between control hCOs (day 150) and prenatal neocortex. *Right: Pearson's correlation* of *NRXN1* exon inclusion levels across splicing sites (SS1–SS6) between control hCOs (day 150) and adult PFC.

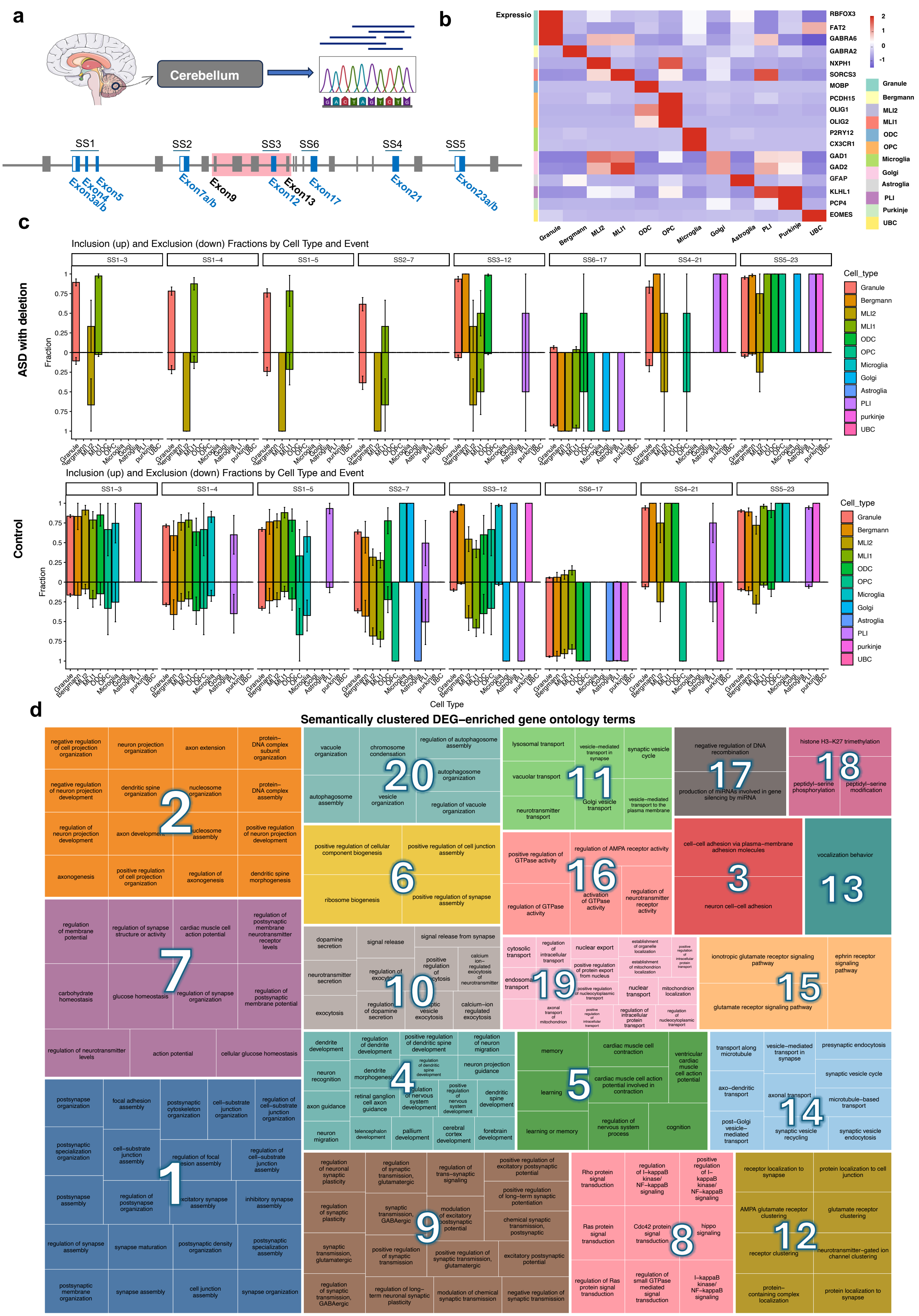
a
Cerebellum
SS1
SS2
SS3
SS6
SS4
SS5
Exon3a/b
Exon4
Exon5
Exon7a/b
Exon9
Exon12
Exon13
Exon17
Exon21
Exon23a/b
b
Expressio
RBFOX3
FAT2
GABRA6
GABRA2
NXPH1
SORCS3
MOBP
PCDH15
OLIG1
OLIG2
P2RY12
CX3CR1
GAD1
GAD2
GFAP
KLHL1
PCP4
EOMES
2
1
0
-1
Granule
Bergmann
MLI2
MLI1
ODC
OPC
Microglia
Golgi
Astroglia
PLI
Purkinje
UBC
c
Inclusion (up) and Exclusion (down) Fractions by Cell Type and Event
ASD with deletion
Fraction
SS1–3
SS1–4
SS1–5
SS2–7
SS3–12
SS6–17
SS4–21
SS5–23
Cell_type
Granule
Bergmann
MLI2
MLI1
ODC
OPC
Microglia
Golgi
Astroglia
PLI
purkinje
UBC
Inclusion (up) and Exclusion (down) Fractions by Cell Type and Event
Control
Fraction
Cell Type
d
Semantically clustered DEG–enriched gene ontology terms
negative regulation of cell projection organization
neuron projection organization
axon extension
protein–DNA complex subunit organization
negative regulation of neuron projection development
dendritic spine organization
nucleosome organization
protein–DNA complex assembly
regulation of neuron projection development
axon development
nucleosome assembly
positive regulation of neuron projection development
axonogenesis
positive regulation of cell projection organization
regulation of axonogenesis
dendritic spine morphogenesis
vacuole organization
chromosome condensation
regulation of autophagosome assembly
autophagosome organization
autophagosome assembly
vesicle organization
regulation of vacuole organization
lysosomal transport
vesicle–mediated transport in synapse
synaptic vesicle cycle
vacuolar transport
neurotransmitter transport
Golgi vesicle transport
vesicle–mediated transport to the plasma membrane
negative regulation of DNA recombination
production of miRNAs involved in gene silencing by miRNA
histone H3–K27 trimethylation
peptidyl–serine phosphorylation
peptidyl–serine modification
regulation of membrane potential
regulation of synapse structure or activity
cardiac muscle cell action potential
regulation of postsynaptic membrane neurotransmitter receptor levels
carbohydrate homeostasis
glucose homeostasis
regulation of synapse organization
regulation of postsynaptic membrane potential
regulation of neurotransmitter levels
action potential
cellular glucose homeostasis
positive regulation of cellular component biogenesis
positive regulation of cell junction assembly
ribosome biogenesis
positive regulation of synapse assembly
positive regulation of GTPase activity
regulation of AMPA receptor activity
regulation of GTPase activity
activation of GTPase activity
regulation of neurotransmitter receptor activity
cell–cell adhesion via plasma–membrane adhesion molecules
neuron cell–cell adhesion
vocalization behavior
dopamine secretion
signal release
signal release from synapse
neurotransmitter secretion
positive regulation of exocytosis
calcium ion–regulated exocytosis of neurotransmitter
regulation of exocytosis
exocytosis
regulation of dopamine secretion
synaptic vesicle exocytosis
calcium–ion regulated exocytosis
cytosolic transport
regulation of intracellular transport
nuclear export
establishment of organelle localization
endosomal transport
positive regulation of protein export from nucleus
establishment of mitochondrion localization
nuclear transport
mitochondrion localization
axonal transport of mitochondrion
regulation of intracellular protein transport
ionotropic glutamate receptor signaling pathway
ephrin receptor signaling pathway
glutamate receptor signaling pathway
dendrite development
regulation of dendrite development
positive regulation of dendritic spine development
regulation of neuron migration
neuron recognition
dendrite morphogenesis
neuron projection guidance
axon guidance
retinal ganglion cell axon guidance
regulation of nervous system development
dendritic spine development
neuron migration
telencephalon development
pallium development
cerebral cortex development
forebrain development
memory
cardiac muscle cell contraction
ventricular cardiac muscle cell action potential
learning
cardiac muscle cell action potential involved in contraction
learning or memory
regulation of nervous system process
cognition
transport along microtubule
vesicle–mediated transport in synapse
presynaptic endocytosis
synaptic vesicle cycle
axo–dendritic transport
axonal transport
microtubule–based transport
post–Golgi vesicle–mediated transport
synaptic vesicle recycling
synaptic vesicle endocytosis
postsynapse organization
focal adhesion assembly
postsynaptic cytoskeleton organization
cell–substrate junction organization
regulation of cell–substrate junction organization
postsynaptic specialization organization
cell–substrate junction assembly
regulation of focal adhesion assembly
regulation of cell–substrate junction assembly
postsynapse assembly
regulation of postsynapse organization
excitatory synapse assembly
inhibitory synapse assembly
regulation of synapse assembly
synapse maturation
postsynaptic density organization
postsynaptic specialization assembly
postsynaptic membrane organization
synapse assembly
cell junction assembly
synapse organization
regulation of neuronal synaptic plasticity
regulation of synaptic transmission, glutamatergic
regulation of trans–synaptic signaling
positive regulation of excitatory postsynaptic potential
regulation of synaptic plasticity
synaptic transmission, GABAergic
modulation of excitatory postsynaptic potential
positive regulation of long–term synaptic potentiation
chemical synaptic transmission, postsynaptic
synaptic transmission, glutamatergic
positive regulation of synaptic transmission
positive regulation of synaptic transmission, glutamatergic
excitatory postsynaptic potential
regulation of synaptic transmission, GABAergic
regulation of long–term neuronal synaptic plasticity
modulation of chemical synaptic transmission
negative regulation of synaptic transmission
Rho protein signal transduction
regulation of I–kappaB kinase/NF–kappaB signaling
positive regulation of I–kappaB kinase/NF–kappaB signaling
Ras protein signal transduction
Cdc42 protein signal transduction
hippo signaling
regulation of Ras protein signal transduction
regulation of small GTPase mediated signal transduction
I–kappaB kinase/NF–kappaB signaling
receptor localization to synapse
protein localization to cell junction
AMPA glutamate receptor clustering
glutamate receptor clustering
receptor clustering
neurotransmitter–gated ion channel clustering
protein–containing complex localization
protein localization to synapse

**Extended Figure 5.**

(a)A schematic of *NRXN1* gene structure as a splicegraph, denoting splice sites (AS1-6), with red shades corresponding on exnoic deletion (exon9-13) genotype.
(b) Heatmap of marker gene expression used for cell-type annotation. Each row represents a marker gene and each column a cell type; darker red indicates higher expression (**Supplementary Table 5)**.
(c) Normalized *NRXN1* splicing events at SS1–SS6 across cell types in control (upper panel) and NRXN1-del cerebellum (lower pannel). Exon inclusion is shown upward ("inclusion") and exclusion downward ("exclusion"). Data represent mean ± SEM.
(d) Treemap showing the top 20 most prevalent biological themes identified from cell type–specific DEGs. Enriched pathways were determined by gene set enrichment analysis and grouped through semantic clustering of Gene Ontology (GO) terms (**Supplementary Table 6**). Biological themes are labeled by their most significant GO term, with numbers indicating prevalence rank, defined by the number of cell types in which each theme is enriched: 1-synapse organization; 2- regulation of neuron projection development; 3 - cell-cell adhesion via plasma-membrane adhesion molecules; 4 - dendritic spine development; 5 - learning or memory; 6 - positive regulation of synapse assembly; 7 - regulation of membrane potential; 8 - regulation of small GTPase mediated signal transduction; 9 - regulation of trans-synaptic signaling; 10 - neurotransmitter secretion; 11 - neurotransmitter transport; 12 - receptor localization to synapse; 13 - vocalization behavior; 14 - vesicle-mediated transport in synapse; 15 - glutamate receptor signaling pathway; 16 - regulation of AMPA receptor activity; 17 - negative regulation of DNA recombination; 18 - peptidyl-serine modification; 19 - positive regulation of intracellular transport; 20 - vacuole organization. Detailed Go terms are listed in **Supplementary Table 7**.

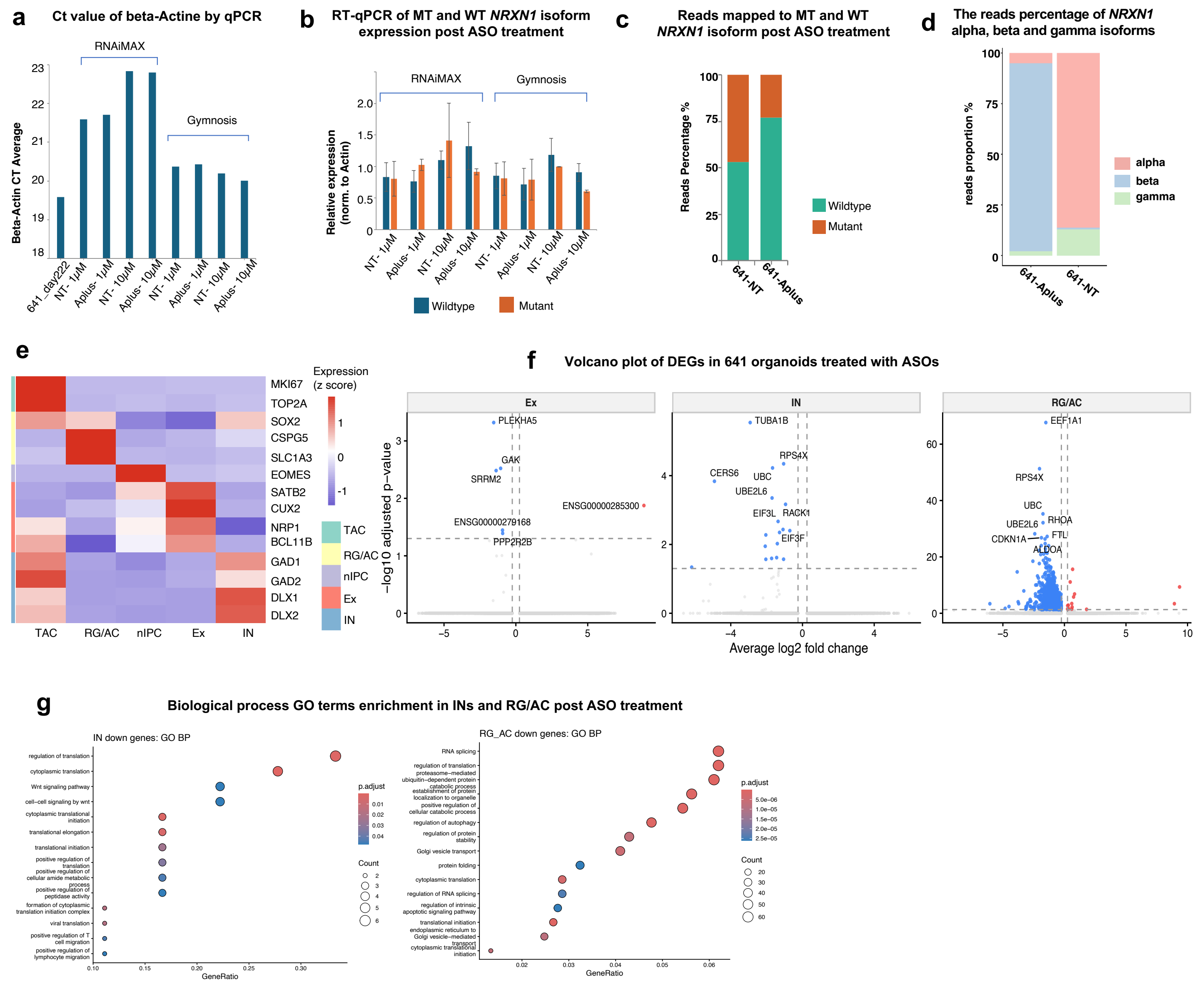

**Extended Figure 6.**

(a)The beta-actin Ct value by qPCR to determine the potential toxicity for organoids treated with 1uM or 10 uM ASO, delivered by RNAiMAX or Gymnosis. The beta -actin Ct value of organoids at day 222 without treatment was used for control.

(b) RT-qPCR of MT and WT *NRXN1* isoform expression post ASO treatment. The organoids were treated with 1uM or 10 uM ASO, delivered by RNAiMAX or Gymnosis. The relative expression level was normalized to beta-actin Ct value.

(c) Percentage of wildtype or mutant *NRXN1* isoforms detected by long-read RACE-seq from 641 hCOs post ASO-treatment.

(d) Percentage of NRXN1 α, β, and γ isoform reads in 641 hCOs treated with ASO-Aplus or ASO-NT, as measured by long-read Capture-seq. A significant enrichment of the β isoform is observed in the 641-Aplus condition ($p < 0.001$, Fisher's exact test).

(e) Heatmap of marker genes expression used for cell-type annotation. Each row represents a marker gene and each column a cell class; darker red indicates higher expression.

(f) Volcano plot for DEGs detected after ASO treatment in excitatory neurons (Ex; left), inhibitory neurons (IN; middle) and RG/AC (right) cells.

(g) Biological process GO terms enrichment in INs and RG/AC post ASO treatment.
